# Spergulin-A strives intra-macrophage leishmanicidal activity by regulating the host P2X_7_R-P_38_MAPK axis

**DOI:** 10.1101/2019.12.14.876391

**Authors:** Niladri Mukherjee, Saswati Banerjee, Sk. Abdul Amin, Tarun Jha, Sriparna Datta, Krishna Das Saha

## Abstract

Current drugs are inadequate for the treatment of visceral leishmaniasis an immunosuppressive ailment caused by *Leishmania donovani*. Regrettably, there is no plant-origin antileishmanial drug present. P2X_7_R is constitutively present on macrophage surfaces and can be a putative therapeutic target in intra-macrophage pathogens with function attributes towards inflammation, host cell apoptosis, altered redox, and phagolysosomal maturation by activating p_38_MAPK. Here we demonstrated that the initial interaction of Spergulin-A (SpA), a triterpenoid saponin with RAW 264.7 macrophages was mediated through P2X_7_R involving the signaling cascade intermediates Ca^++^, P_38_MAPK, and NF-κβ. P_38_MAPK involvement is shown to have specific and firm importance in leishmanial killing with increased NF-κBp65. Phago-lysosomal maturation by Sp A also campaigns for another contribution of P2X_7_R. *In vivo* evaluation of the anti-leishmanial activity of Sp A was monitored through expression analyses of P2X_7_R, P_38_MAPK, and NF-κβ in murine spleen and bone-marrow macrophages and advocated Sp A of being a natural compound of leishmanicidal functions which acted through the P2X_7_R-P_38_MAPK axis.

**SIGNIFICANCE OR IMPORTANCE:** Preciously, this manuscript demonstrated previously unreported initial interaction of Spergulin-A, a triterpenoid saponin isolated from *Glinus oppositifolius* with macrophages through P2X_7_R involving the signaling cascade intermediates Ca^++^, P_38_MAPK, and NF-κβ. Signaling interaction is shown to have specific importance in the leishmanial killing. Phago-lysosomal maturation also campaigns for another contribution of P2X_7_R. *In vivo* evaluation was monitored through P2X_7_R, P_38_MAPK, and NF-κβ in murine spleen and bone-marrow macrophages and advocated Sp A of being a natural compound of leishmanicidal functions which acted through the P2X_7_R-*P_38_MAPK* axis. The result supports that Spergulin-A can provide new lead molecules for the development of alternative drugs against VL. We feel very strongly that this work can be very interesting as it describes a detailed evaluation of leishmanicidal effect by Sp A and thus has every potential to attract a lot of workers especially in the fields of pharmacology, drug development, immunology, as well as parasitology.

## 1. Introduction

The obligate intra-macrophage protozoan parasite *Leishmania donovani* is the causative agent of visceral leishmaniasis (VL). VL is the most severe form of leishmaniasis and if left untreated can reach casualty as high as 100% within 2 years of infection in developing countries (1). VL is widespread in tropical and temperate areas and is endemic in 100 countries with an approximate yearly occurrence of 500,000 cases, of which 70,000 people expire every year (2). Miltefosine, Sodium antimony gluconate, Pentamidine, and Amphotericin B are applied as the choice of chemotherapeutics, however, with reported instances of mild to severe toxicity and resistance (3). *Leishmania* parasites adapted various successful strategies to manipulate different parental activities of macrophage (MΦ) harmful to them, i.e., like antigen presentation, generation of nitric oxide and reactive oxygen intermediates and cytokine production, etc. (4). Despite elaborate reports on leishmanial host-pathogen interaction, explicit revelation of natural compound mediated distortion of signaling cascades to eradicate intracellular leishmanial parasites is rare. Elaboration of plant secondary metabolites that can destabilize parasite propagation and multiplication in addition to direct leishmanicidal activities can advise new therapeutic stratagems.

Botanicals or phytochemicals are widely successful as potent therapeutics against different and various types of human and animal ailments. Nevertheless, unfortunately, VL remains an exception even after serious efforts from different groups of scientists and drug companies (5). Reports of potent plant-derived anti-leishmanial agents are there in the literature (6), however, mostly performed against the vector-borne promastigote form of the parasite (7). Our previous effort demonstrated that triterpenoid saponin, Spergulin-A(Sp A), isolated from a perennial shrub *Glinus oppositifolius* is a potent intracellular antileishmanial compound besides being an immune-modulator (5). It was also reported that triterpenoid saponin of *Careya arborea* and some water-insoluble saponin acted as potent leishmanicidal agents (8–10).

P2X_7_R (a purinergic receptor) is constitutively present on macrophage surfaces and linked with inflammation, apoptosis, generation of Reactive oxygen species (ROS) and nitric oxide (NO), phagolysosomal maturation, and therefore contribute towards intracellular pathogen control (11). Modulation of macrophage biology concerning P2X_7_R expression alteration was monitored in *Leishmania* and other intracellular pathogens like *Toxoplasma*, *Brucella, Cryptosporidium,* and *Microsporidia* (12–16). Expectedly, the absence or underexpression of the P2X_7_R has been linked with under achieved intracellular pathogen control (11). It was also reported that activation of P2X_7_R can trigger mitogen-activated protein kinases (MAPKs), including P_38_MAPKs (17).

Activation of p_38_MAPK in respect to the P2X_7_R over-expression leads to the generation of ROS and NO through nicotinamide adenine dinucleotide phosphate-reduced (NADPH) oxidase (18, 19) and hypothetically can control intra-MФ pathogens including leishmanial parasites. Regulation of inducible nitric oxide synthase (iNOS) expression in macrophages is managed by MAPK signaling molecules with closely associated nuclear factor kappa B (NF-κB) activation (20). It was reported that P_38_MAPK can phosphorylate Rab5 an important molecule during endosome maturation and intracellular leishmanial killing (21). In a previous report of our group, we had elaborated that P_38_MAPK phosphorylation can regulate the maturation of phagolysosomal complexes and subsequent leishmanicidal functions (2). Viscerotropic *L. donovani* parasites can alleviate MAPK phosphorylation and thus neutralize MAPK regulated anti-leishmanial signaling cascades in naive macrophages (22) with controlling NF-kB, the prime transcription factor in NO production (23). P_38_MAPK activation is crucial in *L. donovani* infection abolition is known and related (24). Phagosome-lysosome fusion is maintained by sequential acquisition and removal of different proteins of hydrolases and peptidases and aided in degrading intracellular pathogens like *L. donovani* (25). The phagolysosome fusion is also critically acquainted by enhanced P2X_7_R expression and managed through bivalent calcium ion efflux and RhoA-Phospholipase D controlled pathway (11).

Screening compounds toward purinergic receptor modifications have identified over two thousand structurally diversified compounds and almost one-third of them are natural products (26). For example, teniposide, a semi-synthetic podophyllotoxin derivative can inhibit P2X_7_R, whereas, agelasine and garcinolic acid (26), potentially activate the same. In recent years efforts are made towards the identification of compounds that can modulate P2X receptors and can be potential chemotherapeutics against inflammatory diseases (27, 28). Extracts of the traditional Chinese medicinal plant, *Panax ginseng* can activate P2X_7_ (29) which further identified ginsenosides which are triterpenoid saponins in chemical properties also an activator ofP2X_7_R (30). Conversely, the *R. longifolia* extract and fractions inhibit P2X_7_R leading to their anti-inflammatory function (31). Additionally, there are ample reports of triterpenoid saponins interacting with different groups of MAPKs including P_38_MAPK in conferring their cytotoxic or protective roles (32).

To explore the molecular insights in depicting the intracellular leishmanicidal function of the triterpenoid saponin Sp A, we have emphasized the involvement of P2X_7_R and downstream pathways. We had checked the *in-silico* binding affinity of Sp A with P2X_7_R. P2X_7_R blockage having a hindrance towards the reduction of Sp A mediated leishmanial killing strongly advocates for its association and significance. The involvement of P_38_MAPK is shown to have specific and harsh importance in leishmanial killing as with increased NF-κBp65 mediated ROS and NO production and can be established by simultaneous expression analysis of P_38_MAPK and a glycosylated 91-kDa glycoprotein (Gp91-phox), an NADPH oxidase family member. Phago-lysosomal maturation promotion in mediating intracellular leishmanial killing by Sp A also campaigns for another contribution of P2X_7_R. Finally, the *in vitro* observation was further clarified *in vivo* to conclude Sp A of being a natural compound of leishmanicidal functions acted through the P2X_7_R-P_38_MAPK axis.

## 2. Results

### 2.1. Involvement of the P2X_7_ receptor for the intracellular anti-leishmanial activity of Sp A

At the commencement of the evaluation of antileishmanial affectivity of Sp A against RAW 264.7 MΦ internalized parasites, increasing doses of Sp A (Fig. 1**a**) were applied for 24 h. Sp A was found to be execute intercellular parasites in a dose-dependent manner; however above 30µg/mL of doses, the antileishmanial activity reached a plateau. Therefore, we had selected the dose of 30µg/mL for the detailed assessment of the intracellular anti-leishmanial activity of Sp A. In our previous report, we had elaborated this dose is perfectly safe against the host RAW 264.7 MΦs and that Sp A was not effective against the acellular promastigote or intermediate axenic amastigote forms of *L. Donovani* (5).

**Fig. 1.**
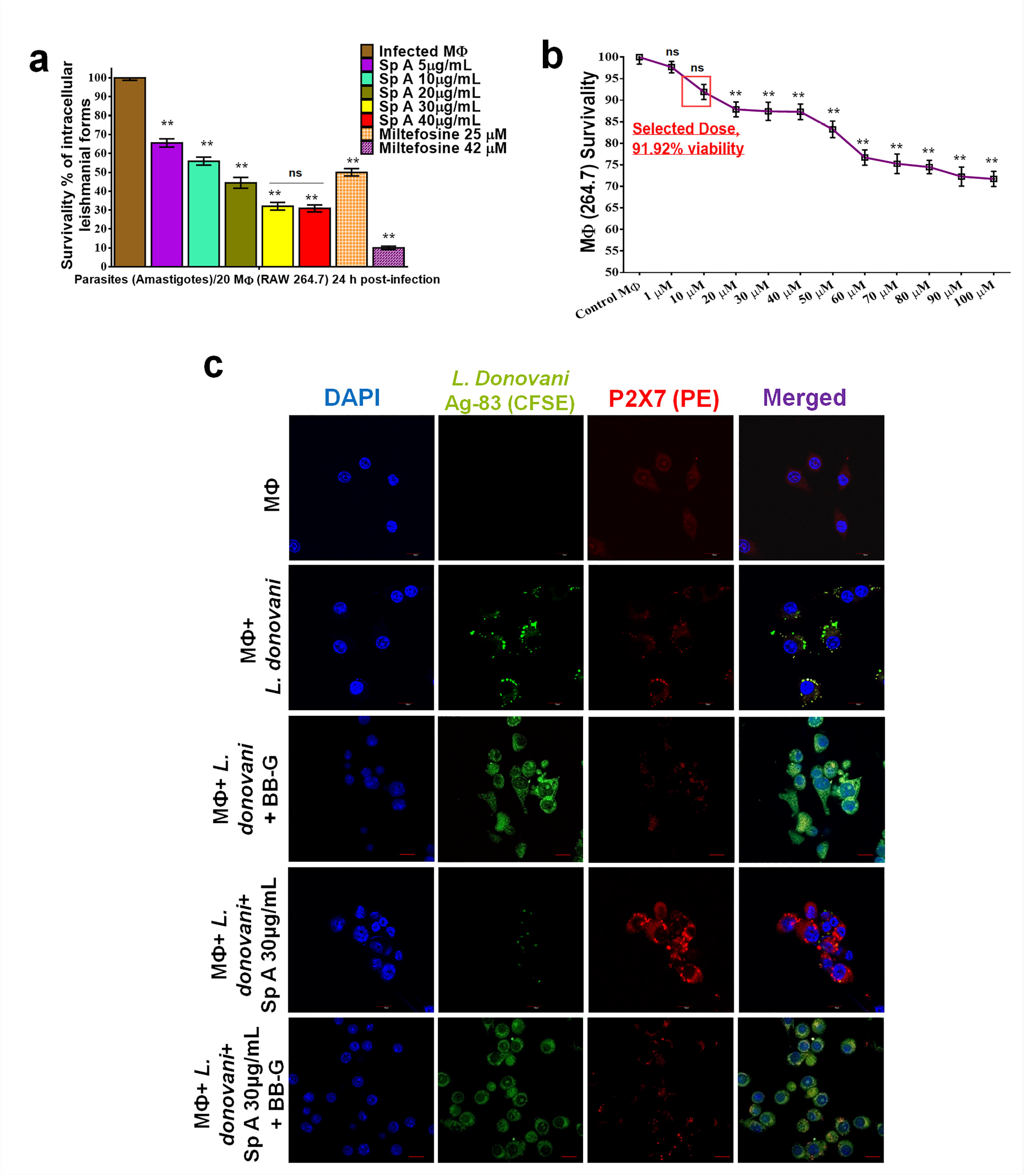
Evaluation of P2X_7_ receptor involvement in the intracellular (RAW 264.7 MΦs) anti-leishmanial activity of Spergulin-A (Sp A). (**a**) Sp A reduces intracellular *L. donovani* expressed as the number of parasites/20 RAW 264.7MΦs dose-dependently after 24 h of treatment with miltefosine as positive control expressing its LC_50_ and LC_90_ values. (**b**) RAW 264.7 MΦ survivability following the P2X_7_ receptor antagonist BB-G application after 24 h. (**c**) Confocal micrographs of consequent expression of CFSE-tagged intracellular *L. donovani* with P2X_7_R expression during Sp A (30µg/mL) and BB-G (10μM) application. Sp A enhanced P2X_7_R expression and intracellular *L. Donovani* removal, whereas, BB-G caused the reversal of the Sp A mediated intracellular parasite removal and P2X_7_R over-expression. Scale bars, 10 μm. Images are representative of three separate experiments. **(a) and (b)**; all values are expressed as mean ± SEM from triplicate assays from three independent experiments; *P*-values ≤0.05 (*) or ≤0.01 (**) vs infected panel except for (b), where the reference is the RAW 264.7 MΦs; ns denotes not significant.

The use of Brilliant Blue G (BB-G) as a selective P2X_7_R antagonist (33) was reported both *in vivo* (34) and *in vitro* systems (35). Different IC50 values for BB-G as a P2X_7_R antagonist had been mentioned in the literature for different cell lines that include a related mouse macrophage cell line (J774.G8) within a dose range of 10μM, (35, 36) however not conclusively against RAW 264.7 MΦs. We evaluated the dose range of BB-G for RAW 264.7 MΦ survival (Fig. 1b). The dose 10μM of BB-G was selected as P2X_7_R antagonist in this study to be applied *in vitro* based upon the high survivability (about 92%) of the exposed cells after 24 h (Fig. 1**b**).

Thereafter, the contribution of P2X_7_R in intracellular *L. donovani* infection and Sp A mediated leishmanicidal function was thoroughly examined (**Supplementary Figure** S1A, S1B and Fig. 1**c**). The presence of Carboxyfluoresceinsuccinimidyl ester (CFSE)-tagged *L. donovani* and subsequent P2X_7_R expression was monitored by FACS during infection and Sp A (30µg/mL) treatment, with or without BB-G, (**Supplementary Figure** S1A i-vi). P2X_7_R expression was highest in the treated panel with the expected lowest CFSE-tagged *L. donovani* existence (**Supplementary Figure** S1A ii, v, vi) and application of BB-G in the treatment panel lowered both the P2X_7_R expression and *L. donovani* removal (**Supplementary Figure** S1A iii-vi). Sp A treatment exhibited a significant rise of P2X_7_R expressing RAW 264.7 MΦs of about 44.3% of the parent population in connection with intracellular parasite removal. Whereas, in all other panels P2X_7_R expressing RAW 264.7 MΦs were about or below 5% of the parent population with or without BB-G application (**Supplementary Figure** S1A i-vi). It was also noticed that BB-G application significantly lowered Sp A mediated intracellular *L. donovani* removal (**Supplementary Figure** S1A iv-vi) and thus affirm the participation of P2X_7_R in Sp A mediated intracellular leishmanicidal activity.

The dose-dependent implication of Sp A in intracellular leishmanial removal and subsequent aversion by BB-G (P2X_7_R antagonist) application was monitored (**Supplementary Figure** S1B). It was observed that BB-G application lowered the leishmanicidal property of Sp A significantly when applied simultaneously with Sp A. Confocal micrographs of CFSE-tagged intracellular *L. donovani* with subsequent P2X_7_R expression were monitored for Sp A treatment as well as BB-G application (Fig. 1**c**). It was obvious from the micrographs that Sp A treatment was able to increase P2X_7_R expression and intracellular *L. donovani* removal, however, application of BB-G reverses the intracellular leishmanicidal function of Sp A exposure (Fig. 1**c**).

This experimental evidence confirmed that Sp A mediated its leishmanicidal property with close assistance with P2X_7_R and subsequent signaling pathways linked with over-expression of it. Therefore, downstream signaling intermediates associated with P2X_7_R should be closely monitored for accessing the precise involvement of P2X_7_R in Sp A mediated leishmanial killing.

### 2.2. Sp A treatment has effects on the P2X_7_ receptor-dependent intracellular Ca^++^ release

It is documented that P2X_7_R activation can lead to Ca^++^ influx and result in mycobacterial removal (37). This Ca^++^ influx is definitely of great consequence in case of intracellular leishmanial killing as this increased Ca^++^titre is linked with phagolysosomal fusion, P_38_MAPK activation, and increased release of NO and ROS along with the altered release of pro and anti-inflammatory cytokines; all of them have leishmanicidal function actively or passively. Thus, it is important to monitor the Ca^++^ influx during Sp A treatment and also link the event with P2X_7_R activation. A set of FACS analyses with P2X_7_R and Ca^++^ corresponding to Fura2M was represented both graphically and through histograms of fluorescence index and percentage of cells expressing the P2X_7_R and Ca^++^for normal, infected and Sp A treated RAW 264.7 MΦs (Fig. 2**a** and **Supplementary Figure** S2A, S2B). The mean fluorescence index equivalent of the expressions of P2X_7_R and Ca^++^ was significantly higher in Sp A treated panels and that too in a dose-dependent manner compared to either normal or infected MΦs (RAW 264.7) (Fig. 2**a** and **Supplementary Figure** S2A, S2B). However, the application of BB-G lowered both of their expressions compared to the corresponding BB-G unexposed panels (**Supplementary Figure** S2A). Subsequently, the percentage of P2X_7_R and Ca^++^ expressing cells against the parent population was also higher for the treated panels, and the application of BB-G not only lowered the P2X_7_R expressing population but also have diminished the effect on Ca^++^ expressing cells (**Supplementary Figure** S2B). These results strongly advocated the involvement and dependence between P2X_7_R expression and Ca^++^ influx in Sp A mediated anti-leishmanial function.

**Fig. 2.**
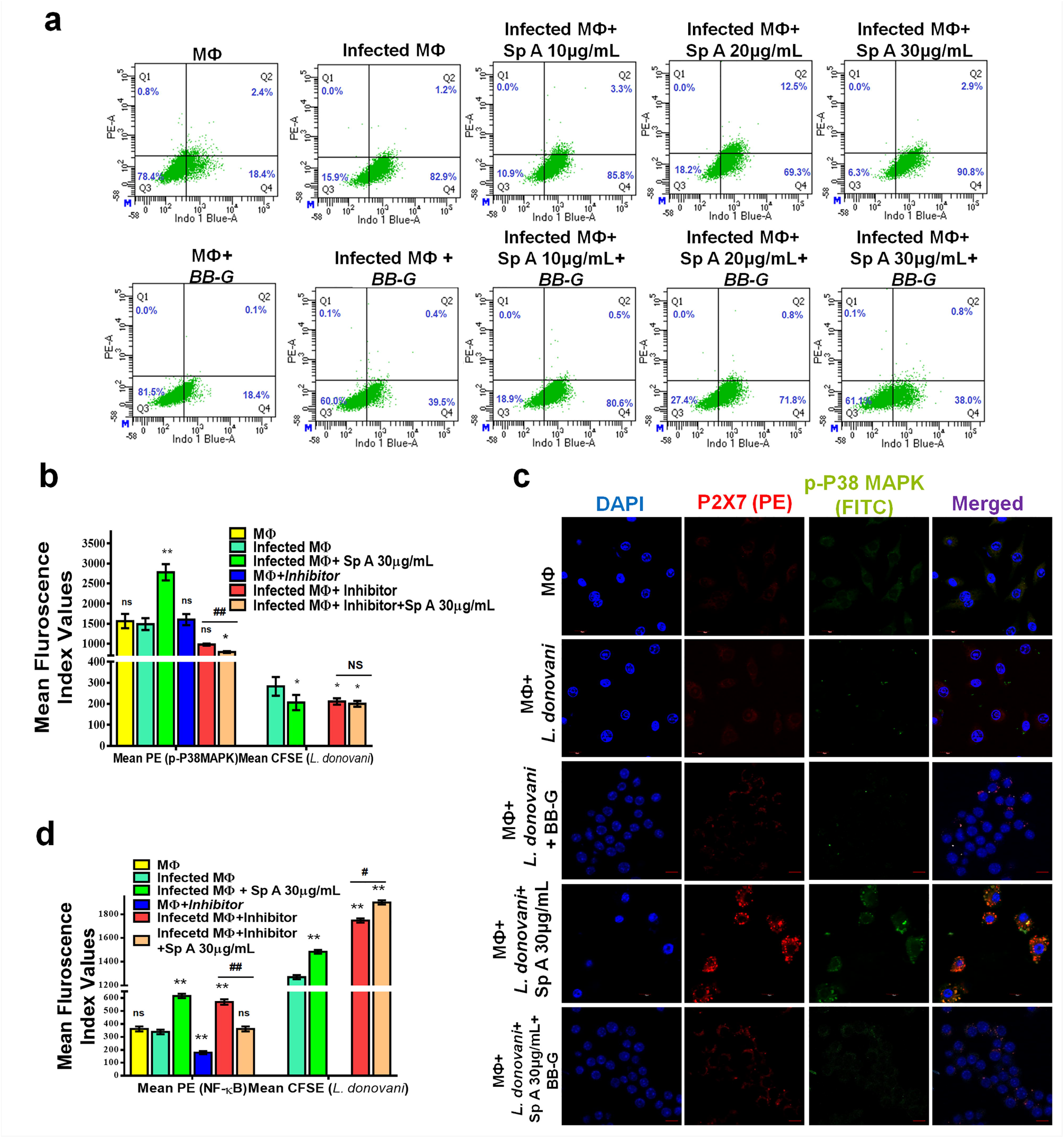
Ca^++^ influx evaluation and P_38_MAPK expression alteration with P2X_7_R expression analyses during Sp A treatment. **(a)** FACS analyses with P2X_7_R expression and Ca^++^ corresponding to Fura2M. The four quadrants represent different stages of P2X_7_R and Ca^++^ expressing populations, i.e., Q1 (only P2X_7_R expressing), Q2 (both P2X_7_R and Ca^++^ expressing), Q3 (double negative), and Q4 (only Ca^++^ expressing). Images are representative of three separate experiments. **(b)** Flow cytometric analyses of p-P_38_MAPK expression alteration towards *L. donovani* infection, Sp A treatment, and P_38_MAPK-inhibitor (SB203580) application and their co-dependence in Sp A mediated anti-leishmanial effect with the histogram of mean fluorescence values of p-P_38_MAPK and CFSE-tagged intracellular (RAW 264.7) *L. donovani* in respective populations. **(c)** Confocal micrographs of co-expression analyses of P2X_7_R and P_38_MAPK in *L. donovani* parasitized, Sp A and BB-G exposed RAW 264.7MΦs. Scale bars, 10 μm. **(d)** Flow cytometric analyses of NF-κBp65 expression alteration towards *L. donovani* infection, Sp A treatment, and NF-κB-inhibitor (Curcumin) application and their co-dependence in Sp A mediated anti-leishmanial effect with the histogram of mean fluorescence values of NF-κBp65 and CFSE-tagged intracellular *L. donovani* in respective populations. **(a) and (c);** images are representative of three separate experiments. **(b) and (d);** all values are expressed as mean ± SEM from triplicate assays from three independent experiments (*P*-values **≤**0.05 (*) or **≤**0.01 (**) vs infected panel).

### 2.3. P_38_MAPK expression alteration affects Sp A mediated anti-leishmanial activity with P2X_7_ receptor involvement

The contribution of P_38_MAPK was evaluated in *L. donovani* infection development and also its expression alteration due to Sp A treatment by flow cytometry (Fig. 2**b** and **Supplementary Figure** S3A). Apart from the panels where P_38_MAPK inhibitor (SB203580) was used (Fig. 2**b** and **Supplementary Figure** S3A iv-vi), almost all the cells found expressing P_38_MAPK were either infected or treated with Sp A (**Supplementary Figure** S3A i-iii). The difference lies in the expression level of P_38_MAPK (Fig. 2**b**). Compared to the control MФ panel, infection with *L. donovani* comprehensively reduced P_38_MAPK (Fig. 2**b** and **Supplementary Figure** S3A ii). However, Sp A treatment increased the P_38_MAPK intensity (Fig. 2**b** and **Supplementary Figure**S3Aiii) and positively correlate with the reduced intracellular parasitic count because when SB203580 was used the parasitic count had decreased along with the P_38_MAPK expression (Fig. 2**b** and **Supplementary Figure**S3A).

Co-expression analyses of P2X_7_R and P_38_MAPK were evaluated by confocal microscopy (Fig. 2**c**). For the control RAW 264.7 MΦs, both the expressions of P2X_7_R and P_38_MAPK were at a basal level and for the infected and infected with BB-G panels, their expression got down-regulated. Whereas, in the Sp A treated panel both P2X_7_R and P_38_MAPK were overexpressed several folds (Fig. 2**c**) per its anti-leishmanial effect. Application of BB-G, the P2X_7_R antagonist also lowered the expression of P_38_MAPK. Inhibitor-mediated underexpression of P_38_MAPK in the treated panels demonstrated compromised intracellular parasite removal (Fig. 2**b** and **Supplementary Figure** S3A).

The confocal micrographs (Fig. 2**c**) also suggested that P2X_7_R activation can positively regulate P_38_MAPK expression and both of them are essential in the intracellular Sp A mediated anti-leishmanial effect of which P_38_MAPK act downstream of P2X_7_R.

### 2.4. NF-κBp65 expression alteration in Sp A mediated leishmanicidal activity

Likewise, the connection of NF-κB in Sp A-mediated infection control was also monitored (Fig. 2**d** and **Supplementary Figure** S3B). The trend was almost alike of P_38_MAPK expression, parasitic infection down-regulated the expression of NF-κB, and Sp A treatment up-regulated their expression when applied (Fig. 2**d** and **Supplementary Figure** S3B). The inhibitor was noticed to down-regulate NF-κB expressions with a moderate increase of the parasitic load even when co-applied with Sp A (Fig. 2**d** and **Supplementary Figure** S3B vi).

### 2.5. ROS and NO participate in anti-leishmanial effects of Sp A

Changes in the production of ROS, as well as NO in respect to *L. donovani* infection and Sp A treatment, were measured by flow cytometry (**Supplementary Figure** S4A, S4B) and confocal micrographs (Fig. 3**a**) with the aid of fluorescent dyes dichlorodihydrofluorescein diacetate (DCFDA) and Diaminofluorescein-2 diacetate (DAF-2 DA) respectively. It was observed that ROS production was downregulated for *L. donovani* infection. In comparison to infected MФs, Sp A (30μg/mL) exposure elevated ROS production in the infected MФs (**Supplementary Figure** S4A iv, vi) which is in the alignment of the lessened parasite load in them (Fig. 3**a**). However, when N-acetylcysteine (NAC) has applied the ROS abrogation (**Supplementary Figure** S4A v, vi) also leads to substantial loss of the parasiticidal activity of Sp A (Fig. 3**a**).

**Fig. 3.**
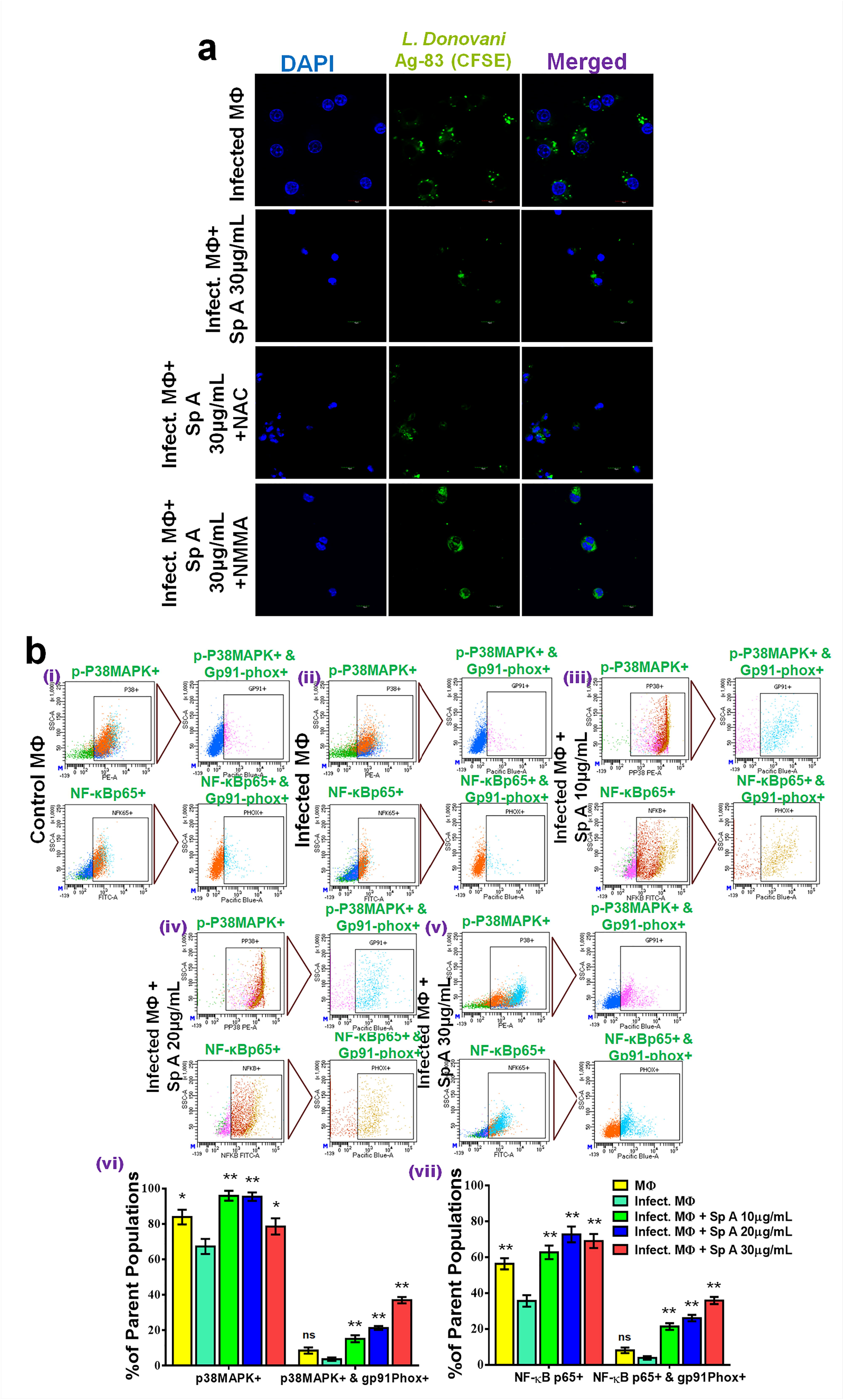
**(a)** Changes in CFSE-tagged intracellular (RAW 264.7MΦs) *L. donovani* parasites were micrographed for Sp A treatment and co-application of NAC and NMMA for evaluation of ROS and NO involvement in Sp A mediated anti-leishmanial effect by confocal microscopy, scale bars, 10 μm. Images are representative of three separate experiments. **(b)** Co-expression alteration of p-P_38_MAPK and Gp91-phox, p-NF-κBp65, and Gp91-phox in Sp A mediated anti-leishmanial effect. Sp A at different doses (10, 20, and 30μg/mL) applied in the infected RAW 264.7MΦs found to elevate levels of Gp91-phox for p-P_38_MAPK positive RAW 264.7MΦs as evaluated by FACS. Histograms of percent parent populations for p-P_38_MAPK positive MФs and both Gp91-phox and p-P_38_MAPK for different infected and Sp A treated populations denoted their inter-connections. Sp A at different doses (10, 20, and 30μg/mL) applied in the infected RAW 264.7MΦs found to elevate levels of Gp91-phox for NF-κBp65 (Rel A) positive RAW 264.7MΦs as evaluated by FACS. Histograms of percent parent populations for NF-κBp65positive RAW 264.7MΦs and both Gp91-phox and NF-κBp65 for different infected and Sp A treated populations denoted their inter-connections. All values presented in the histograms are expressed as mean ± SEM from triplicate assays from three independent experiments (*P*-values **≤**0.05 (*) or **≤**0.01 (**) vs infected panel).

Sp A (30μg/mL) treatment was found to encourage NO production in the infected MФs (**Supplementary Figure** S4B v, vii). It was found that the NO production has a positive effect in checking the parasite load inside the MФs as N-monomethyl-L-arginine (NMMA) application in the Sp A treated MФs reduced the effectivity of the compound (**Supplementary Figure** S4B vi, S4C and Fig. 3**a**). Most increment (5.6-fold) in extracellular NO was noticed for 30 µg/mL of Sp A treatment, followed by 20 μg/mL (3.8-fold), application NMMA lowered extracellular NO production to 2.4-fold almost like that of Sp A at 10 μg/mL compared to the infected panel, extracellular NO in which found to be the least (**Supplementary Figure** S4C).

### 2.6. Expression alteration of p-P_38_MAPK, p-NF-κBp65, and Gp91-phox in intervening ROS and NO involvement of Sp A mediated anti-leishmanial effect

Gp91-phox the glycosylated subunit of the heterodimer phagocytic NADPH oxidase flavocytochrome b558, plays a crucial role in regulating the redox status of MФs (2) and thus can serve as a marker of the alteration of oxidative stress due to Sp A treatment. Sp A at applied doses (10, 20 and 30μg/mL) in the infected MФs also evaluated for Gp91-phox against both p-P_38_MAPK and NF-κBp65 (Rel A) positive MФs by FACS (Fig. 3**b**) and confocal microscopy (**Supplementary Figure S5**) (for only p-P_38_MAPK). Gp91-phox expression was simultaneously up-regulated with increasing doses of Sp A along with the increment of p-P_38_MAPK expression (Fig. 3**b**i-v (upper panels) and Fig. 3**b**vi). Likewise, Gp91-phox expression was also found to be elevated with respect to NF-κBp65 expression along with Sp A in a dose-dependent manner (Fig. 3**b**i-v (lower panels) and Fig. 3**b** vii).

Confocal images provided evidence that Sp A with the increasing doses had increased p-P_38_MAPK and Gp91-phox expression simultaneously (**Supplementary Figure S5**). Infection with *L. donovani* lowered both p-P_38_MAPK and Gp91-phox expression compared to the control (**Supplementary Figure S5**).

### 2.7. Evaluation of p-P_38_ MAPKagainst total P_38_ MAPK and p-NF-κBp65 against NF-κBp65

Western blot analyses (Fig. 4**a** i, ii) showed that p-P_38_ MAPK presence was highest for Sp A treated (30μg/mL) when normalized against total P_38_MAPK. Both the inhibitors of P_38_ MAPK and P2X_7_R considerably downregulated related phosphorylation of P_38_MAPK (Fig. 4**a**i) and the phosphorylation of P_38_MAPK was found to be the lowest for the infected RAW 264.7 MΦs. While LPS found to phosphorylate NF-κBp65 at a higher amount than control or infected RAW 264.7 MΦs, here also it was noticed that Sp A treatment can form p-NF-κBp65 considerably higher than infected or LPS exposure (Fig. 4**a** ii).

**Fig. 4:**
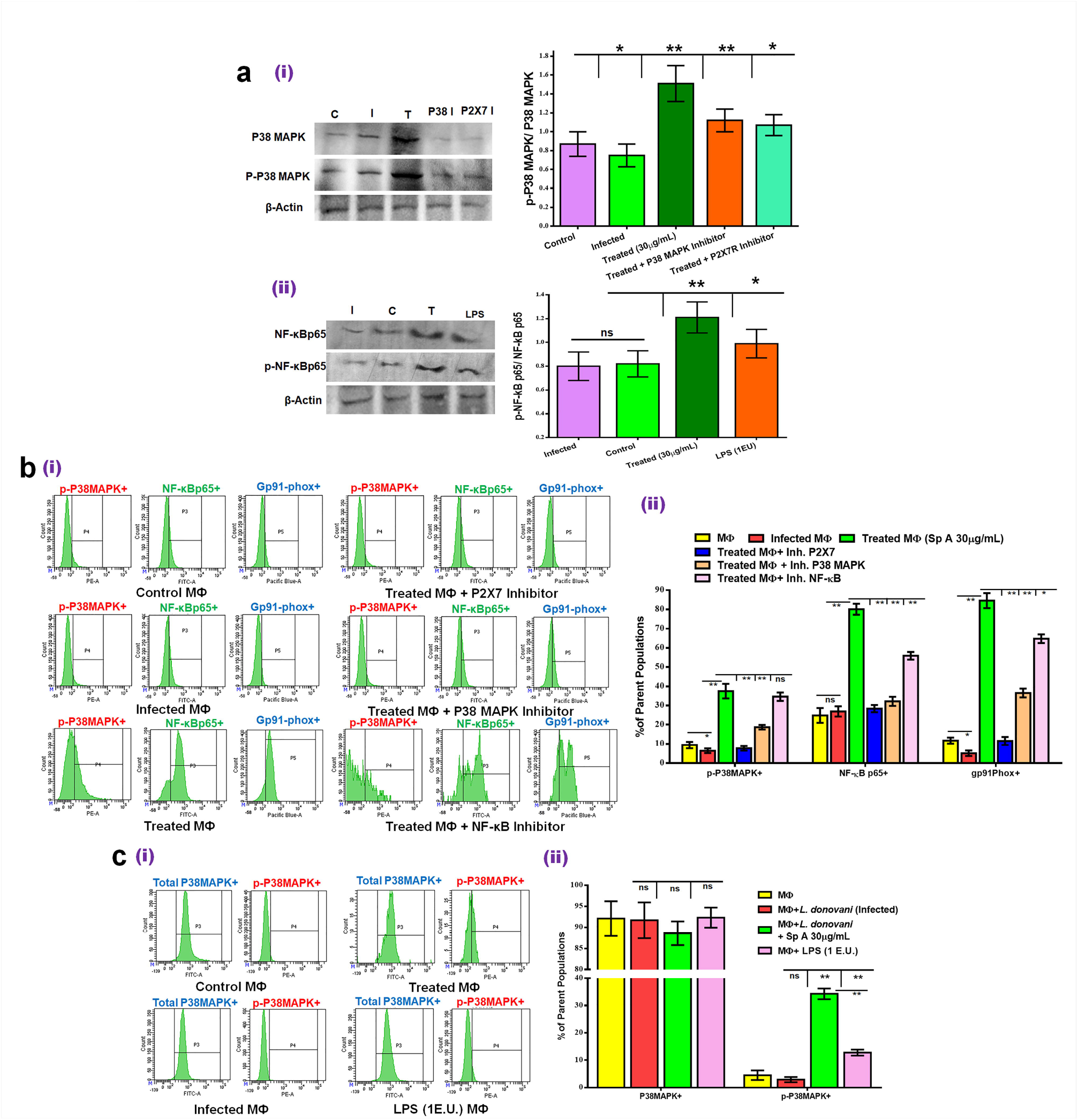
**(a)** Western blot analyses of p-P_38_ MAPK against total P_38_ MAPK and p-NF-κBp65 against NF-κBp65. p-P_38_ MAPK has been normalized against P_38_ MAPK and p-NF-κBp65 was also normalized against NF-κBp65 and simultaneously with the housekeeping protein (β-Actin) expression. (C, control; T, Sp A (30μg/mL treated, I, *L. donovani* infected). **(b)** Evaluation of Sp A activity through P2X_7_R-P_38_MAPK -NF-κB axis and involvement Gp91-phox expression monitored by FACS. **(c)** Confirmation of the relative change in P_38_ MAPK during Sp A treatment. Total P_38_MAPK and p-P_38_MAPK was evaluated by FACS and LPS (1 EU/mL) (EU=endotoxin unit≈ 0.1 to 0.2 ng endotoxin/mL) is used as positive control. Images are representative of three separate experiments. All values presented in the histograms are expressed as mean ± SEM from triplicate assays from three independent experiments (*P*-values **≤**0.05 (*) or **≤**0.01 (**) as indicated in the figure).

### 2.8. Sp A acts through P2X_7_R-P_38_MAPK -NF-κB axis and involvement Gp91-phox expression

FACS analyses were performed to confirm P2X7R-P_38_MAPK -NF-κB axis during Sp A mediated the anti-leishmanial effect of Sp A and simultaneous involvement of Gp91-phox with inhibitors of these key mediators (Fig. 4**b**, Fig. 4**c**). Sp A at 30μg/mL showed upregulated expression and percent population of p-P_38_MAPK, NF-κBp65, and Gp91-phox expressing cells significantly (*P*-values **≤**0.01 (**) higher than that of the infected panel. However, to confirm the axis antagonist or inhibitor of P2X_7_R, P_38_MAPK and NF-κB were added to the treatment panel, and expression of p-P_38_MAPK, NF-κBp65, and Gp91-phox was monitored by FACS (Fig. 4**b**, Fig. 4**c**). This led us to evaluate the P2X7R-P_38_MAPK -NF-κBaxis as well as the involvement of P_38_MAPK and NF-κB in Gp91-phox expression. Application of P2X_7_R antagonist downregulated expression of all the subsequent effectors, p-P_38_MAPK, NF-κBp65, and Gp91-phox like that of the P_38_MAPK inhibitor panel. It was noticed that P2X_7_R acts upstream of P_38_MAPK and is mediated through Ca^++^ release (Fig. 2**a**, Fig. 2**c**). Subsequently, application of the P_38_MAPK inhibitor also downregulated the expression of all of these proteins and at the same time, the NF-κB inhibitor was unsuccessful in significantly downregulating p-P_38_MAPK expression and lowered the expression of NF-κBp65 and Gp91-phox. Thus, from this experiment, it can be stated that both P_38_MAPK and NF-κB can positively upregulate Gp91-phox expression and Sp A mediates its intracellular anti-leishmanial function through P2X7R-P_38_MAPK -NF-κB axis.

### 2.9. Confirmation of the relative change in P_38_ MAPK during Sp A treatment

P_38_MAPK is usually present in an inactive (non-phosphorylated) state in normal, unstimulated RAW 264.7 MΦs used as control and the activation or phosphorylation of this P_38_MAPK to p-P_38_MAPK is necessary for the downstream signaling. Phosphorylation of P_38_MAPKor activation can occur to stimulants such as bacterial lipopolysaccharide (LPS), or application of the therapeutic compound. Hereby the application of Sp A found to be important for the intracellular leishmanial control. Total P_38_MAPK and p-P_38_MAPK was evaluated by FACS and LPS (1 EU/mL) (EU=endotoxin unit≈ 0.1 to 0.2 ng endotoxin/mL) is used as positive control (Fig. 4**c**). *L. donovani* infection or Sp A treatment or LPS exposure was found not to significantly alter the total P_38_MAPK levels across the experimental panels. However, the p-P_38_MAPK level has been significantly (*P*-values ≤0.01 (**) upregulated in the Sp A treated and LPS treated panels (Fig. 4**c**, i, ii), and the phosphorylation is most in the Sp A treated parasitized RAW 264.7 MΦs which is significantly higher than LPS treated one. Thus, it implies that Sp A able to phosphorylate the P_38_MAPK convincingly enough for the subsequent pathway needed for its anti-leishmanial function.

### 2.10. Phago-lysosomal maturation promotion by Sp Awith respect to its anti-leishmanial activity

Phagolysosomal maturation is considered a significant process to eradicate the intracellular pathogens of phagocytic cells and the pathogens promoted multiple ways to evade this sequential process (25). Sp A mediated dynamics of Rab5, an early phagosomal marker, and CathepsinD, a late phagosomal marker was observed by confocal microscopy (Fig. 5**a**). *L. donovani* infection had not mediated any significant visible alterations of these markers compared to the control panel. Whereas, the dose-dependent application of Sp A amplified both Rab5 and CathepsinD expression and documented phagolysosomal fusion during the leishmanicidal property of Sp A (Fig. 5**a**).

**Fig. 5.**
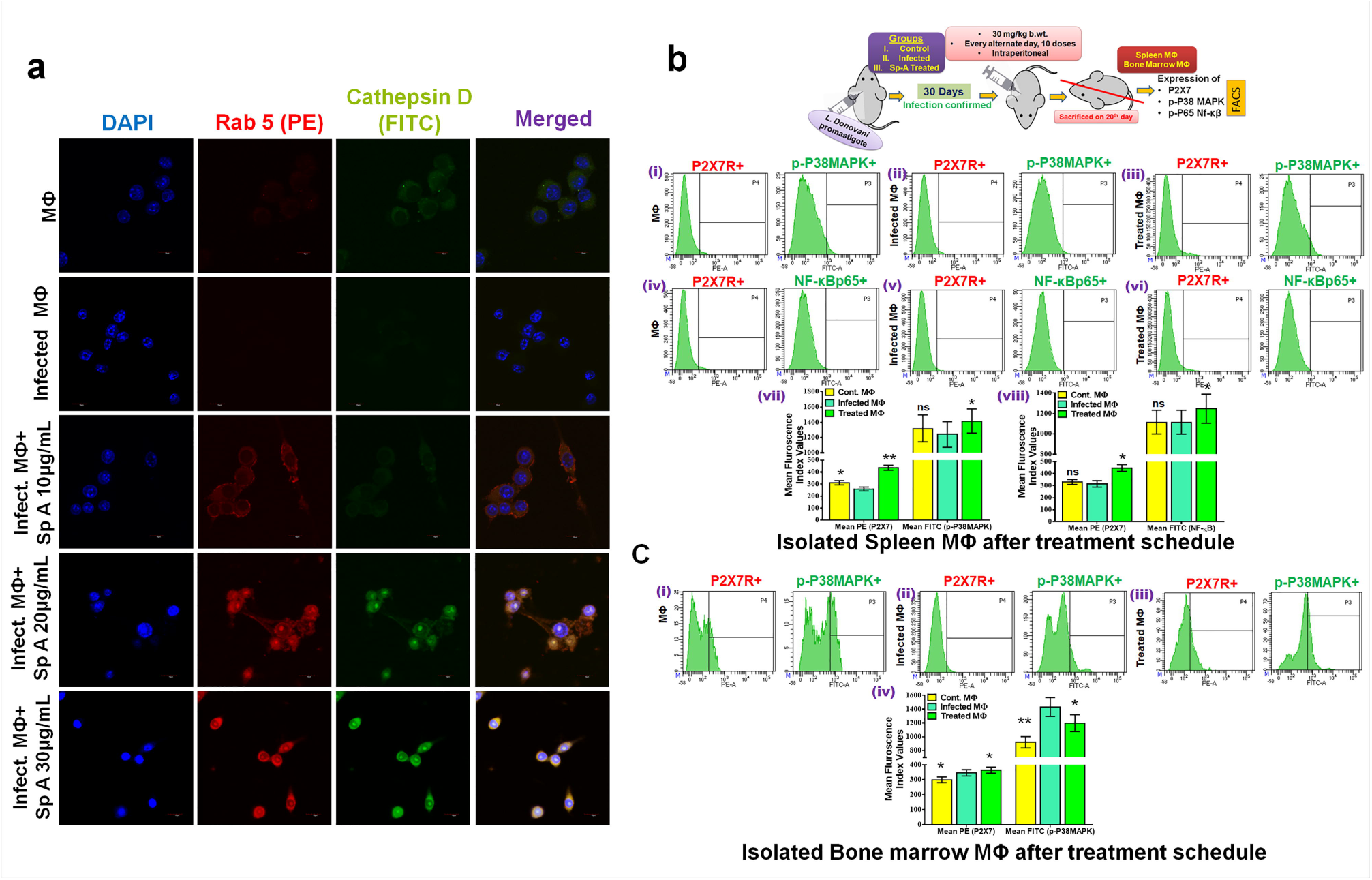
**(a)** Dynamics of phagolysosomal maturation in demonstrating the dose-dependent anti-leishmanial effect of Sp A. Early phagosome marker Rab5 expression got upregulated in the treated panel with respect to the control and infected panels. Cathepsin-D (Late phagolysosomal marker) expression also got upregulated with the exposure of Sp A and signifies the phagolysosomal maturation process. Images are representative of three separate experiments, scale bars, 10 μm. **(b)** *In vivo* evaluation of the anti-leishmanial effect of Sp A with P_38_-MAPK and P2X_7_ receptor involvement.Graphical presentation of the *in vivo* treatment schedule and observation parameters. After confirmation of the *L. donovani* infection in the BALB/C mice after30 days of initial infection, infected mice were divided into two groups, one Sp A treated and other infected control groups arbitrarily with an additional one un-infected control group of 7±2 animals in each group. Un-infected control and treated groups received 30mg/kg b.wt. of Sp A for every alternate day for 10 doses then sacrificed and spleen and bone marrow MΦs were harvested and examined for P2X_7_R, p-P_38_MAPK, and NF-κBp65 expression. Mice from each group were sacrificed and spleen MΦs were harvested for each group and evaluated by flow cytometry for the linked expressions of P2X_7_R-p-P_38_MAPK and P2X_7_R-NFκBp65. Mean fluorescence index values equivalent to the expressions of both P2X_7_R-p-P_38_MAPK and P2X_7_R-NFκBp65from spleen MΦs of different experimental mice groups. **(c)** Mice from each group were sacrificed and bone marrow MΦs were harvested from each group and evaluated by flow cytometry for the linked expressions of P2X_7_R-p-P_38_MAPK. Mean fluorescence index values equivalent of the expressions of both P2X_7_R-p-P_38_MAPK from bone marrow MΦs of different experimental mice groups. Images are representative of three separate experiments. Histogram values (**b** vii, viii, and **c** iv) are expressed as mean ± SEM from triplicate FACS assays (*P*-values **≤**0.05 (*) or **≤**0.01 (**) vs infected control groups, ns denotes not significant).

### 2.11. In vivo evaluation of the anti-leishmanial activity of Sp A in lights of P_38_-MAPK and P2X_7_ receptor involvement

Evaluation of the intracellular anti-leishmanial activity of Sp A *in vitro* against RAW 264.7 MΦs strongly favoured the involvement of P2X_7_R activation and subsequent participation ofP_38_MAPK and NF-κBp65. Therefore, an *in vivo* evaluation of the anti-leishmanial activity of Sp A with a demonstration of the expression alteration of the aforementioned key mediators can link the *in vitro* observations *in vivo*. After the treatment schedule, (Un-infected control and Treated groups received 30mg/kg b.wt. of Sp A, intraperitoneally in an emulsified saline solution of 1 mg/mL concentration for every alternate day for 10 doses, whereas, the infected control group received only the vehicle) mice were sacrificed. Spleen and bone marrow MΦs were harvested for each group and evaluated by flow cytometry for the linked expressions of P2X_7_R-P_38_MAPK and P2X_7_R-NF-κBp65 against spleen MΦs (Fig. 5**b**) and P2X_7_R-P_38_MAPK against bone marrow MΦs (Fig. 5**c**).

From the observations made against spleen MΦs, the treated panels showed elevated P2X_7_R activation against both control and infected panels (Fig. 5**b** vii, viii). Subsequent expressions of P_38_MAPK (Fig. 5**b** vii) and NF-κBp65 (Fig. 5**b** vii) showed the same pattern; the highest for the treated panels. However, for the Bone marrow MΦs, although P2X_7_R expression was highest in the treated panel (Fig. 5**c**iii, iv), the P_38_MAPK expression was less than the infected group (Fig. 5**c** ii, iv).

The *in vivo* findings confirmed the *in vitro* observations and were perfectly align the pathway of action speculated from the *in vitro* findings for the intracellular activity evaluation of Sp A; precisely with the involvement of P2X_7_R, Ca^++^ influx, P_38_MAPK, NF-κBp65, ROS, NO, and phagolysosomal maturation being the key mediators. A graphical presentation of the step-wise anti-leishmanial activity propagation by Sp A has been demonstrated in Fig. 6**d**.

**Fig. 6.**
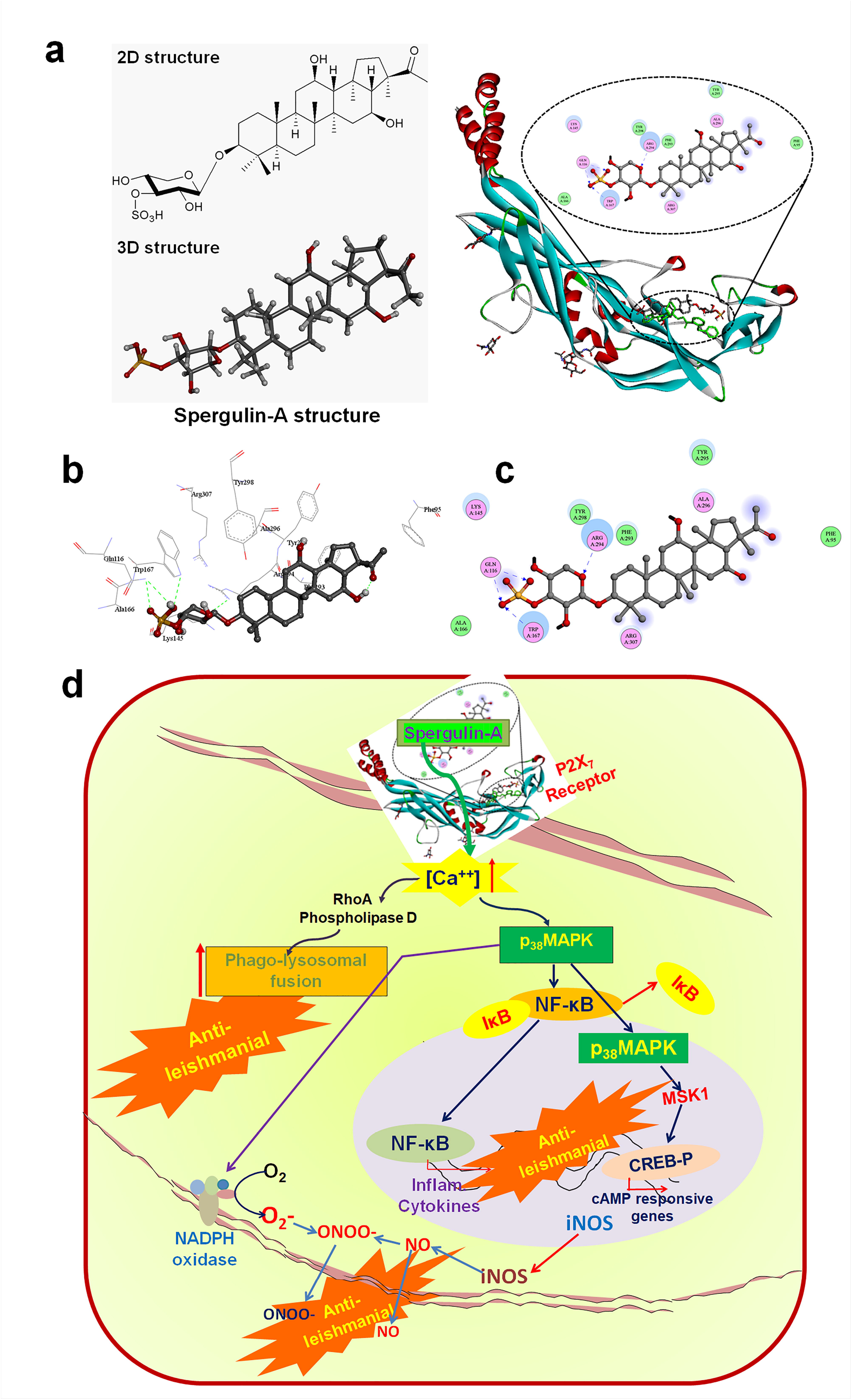
*In silico* docking study of the interdependence of Sp A with P2X7 receptor. **(a)** The proposed interaction investigated compound Sp A with the P2X_7_ receptor. The green-coloured stick is the inbound crystal compound. **(b)** The proposed 3D interaction investigated compound Sp A with the amino acid residues of the P2X_7_ receptor. **(c)** The proposed 2D interaction investigated compound Sp A with the amino acid residues of the P2X_7_ receptor. **(d)** A graphical presentation of the step-wise anti-leishmanial activity propagation by Sp A.

### 2.12. In silico binding assessment of Spergulin A with P2X_7_ Receptor

The possible binding mode of Sp A with its putative target P2X_7_R was analysed by *in silico* methods. A molecular docking study was employed to check the binding mode of the investigated compound i.e., Sp A towards target P2X_7_R (PDB: 5U1X). The molecular docking result illustrates few hydrogen bonding interactions between the Sp A and the P2X_7_ receptor as shown in Fig. 6**a**. The docked conformation of Sp A with P2X_7_R indicates that the sulfonyl function interacts with Gln116, Trp167 (Fig. 6**b**). The amino acid residue Arg295 is found to form another hydrogen bonding interaction with the oxygen atom of the pyran nucleus (Fig. 6**c**). Moreover, it is also noticed that the non-sugar part of Sp A is exposed to the solvent-accessible surface area of the receptor.

## 3. Discussion

Leishmaniasis is a poorly investigated disease mainly affecting people in developing countries with challenged health practices. *Leishmania* living within the harsh environment of phagocytes has developed strategies for prompt physiological adaptation, escape from first-line defence systems, and the ability to inhibit several host cell functions (4). The direct influence of the parasite on the macrophages is linked to the suppressed generation of nitric oxide and oxidative burst and also their inhibitory host immune responses (38, 39). Indirectly, *Leishmania* influences the antigen presentation and T cell tasks in a way that macrophage deactivation guides towards parasite endurance mainly by amendment of host cell signaling of different kinases and phosphatases (40). Our previous effort (5) demonstrated that *G. oppositifolius* and an isolated triterpenoid saponin, Sp A was inconsistent with the direct leishmanicidal effect against promastigotes and axenic amastigotes. However, Sp A was effective against intracellular *L. donovani* virulent strain besides being an immune-modulator measured *in vitro* in RAW 264.7 MФs (5). This observation opens for the scope for precise demonstration of the pathways involved in its intracellular anti-leishmanial effect and involvement and possible activation of P2X_7_R has been considered. P2X_7_R was reported to be constitutively expressed on MΦ surfaces and also reported to be associated with signaling cascades of inflammation, apoptosis, and subsequent intracellular pathogen control (11). Additively, intracellular anti-leishmanial attributes of Sp A in pursuing the involvement of the P2X_7_ purinergic receptor are also hypothesized on the previous reports of compounds screened towards the purinergic receptor, one-third of them being natural products (26).

Reduction of Sp A mediated intra-MΦ leishmanial count was observed to be linked with P2X_7_R activation and this reduction was hindered with P2X_7_R antagonist BB-G application and confirmedP2X_7_R involvement and importance (Fig. 1**a-c** and **Supplementary Figure** S1A, S1B). Existing literature reported the involvement ofP2X_7_R during the removal of intracellular parasites namely *Chlamydia*, *Leishmania*, *Toxoplasma* (41, 42), and *Mycobacterium tuberculosis* by mediating strong pro-inflammatory and cytotoxic activity (43). P2X_7_R activation was found to help reduce *L. amazonensis* parasitic load in macrophages via leukotriene B4 (LTB4) production and connected with lesion size and the parasitic load (44, 45). BB-G is a P2X_7_R reversible antagonist acting at the receptor level with thereby preventing the channel from opening. There are various molecules reported that can inhibit P2X_7_R activity and are majorly classified as orthosteric ligands (bind the receptor within the ATP-binding cavity) and allosteric ligands (bind at other than the ATP-binding cavity). Examples of orthosteric ligands are suramin or suramin-like derivatives, ATP derivatives (TNP-ATP, periodate-oxidized ATP [oATP]), tetrazole derivatives (A438079, A839977), and cyanoguanidine derivatives (A740003, A804598) and examples of allosteric ligands are BB-G, AZD9056, KN-62, AZ-11645373, AZ-10606120, GW791343, GSK314181A, GSK1482160, CE-224,535, AFC-5128, JNJ-479655, and EVT-401 (46, 47). Orthosteric ligands are effective at nanomolar or micromolar concentrations and BB-G can also block P2X1, P2X4, and sodium channels (47, 48). However, a point in support of using BB-G is its availability across various laboratories and relative commercial benefits for being cheap. BB-G application is also depicted in related mouse macrophage cell line (J774.G8) within a dose range of 10μM, (35, 36) however not done conclusively against RAW 264.7 MΦs. Therefore, in the present work, the opportunity is taken to evaluate its effectiveness as an orthosteric receptor antagonist against RAW 264.7 MΦs, and concentration-specific mortality had also been evaluated. Though more experiments are needed for complete validation, BB-G emerged as a safe P2X_7_R antagonist against the cell line within the applied dose (10µM) (Fig. 1**b**). Its importance also magnifies on the *in silico* experimental evidence of binding of Sp A with P2X_7_R. Application of BB-G here has reduced Sp A-mediated increase in P2X_7_R expression and therefore suggested that the transcriptional levels of P2X_7_ are affected. P2X_7_Ractivation with extracellular stimuli like ATP can initiate the opening of cation-specific ion channel and result in the influx of Ca^++^ and the efflux of K^+^ and with extended exposure able to create pores in the plasma membrane resulting in the passage of larger molecules (11). Our experimental verification confirmed Sp A activated P2X_7_R mediated Ca^++^ influx (Fig. 2**a** and **Supplementary Figure** S2A, B) and unleashed a key mediator in intracellular leishmanial control. There is an interesting observation during quantitative FACS estimation of P2X_7_R (**Supplementary Figure** S2B). The mean P2X_7_R fluorescence intensity is expectedly higher for 30µg/mL of Sp A treatment, however, the percentage RAW 264.7 MΦs expressing P2X_7_R is higher in 20µg/mL. This is most possibly because compared to 20µg/mL, Sp A at 30µg/mL able to clear more macrophages from the intracellular *L. donovani* and those parasite-cleared cells might not express P2X_7_R and possibly resulted in a lower percentage of the total population expressing P2X_7_R, however, with higher fluorescence intensity for being actively involved in removing the rest of the parasites.

Changes in the concentration of intracellular Ca^2+^ and PI3K activity are essential for proper phagosomal maturation (49). Ca^++^ is a key second messenger and cytosolic build-up of Ca^++^ can lead to synaptic transmission, macrophage activation, and apoptosis (50). More interestingly, the cytosolic Ca^++^ concentration affects the phagosomal maturation process by modulating membrane fusion between phagosomes and lysosomal vesicles by activating phospholipase D via RhoA, important in intracellular *Leishmania* killing (51). Phagolysosomal fusion can also be mediated by regulating the activities of two Ca^2+^-dependent effector proteins, calmodulin and the multifunctional serine/threonine-protein kinase CaMKII (52). Sp A mediated phagolysosomal maturation was confirmed in the treated parasitized RAW 264.7 MΦs (Fig. 5**a**) by examining co-expression of early Rab5 and late CathepsinD phagolysosomal markers in the treated cells and confirmed the event in intracellular parasite control.

An influx of Ca^++^can activate the P_38_MAPK, which in turn stimulates a number of downstream effectors signaling cascades (53). The connection of phosphorylated P_38_MAPK and leishmanial removal was closely monitored with Sp A treatment (Fig. 2**b**,**c**, Fig. 4**c** and **Supplementary Figure** S3A). With infection p-P_38_MAPK level lowered and treatment with Sp A resulted in elevated p-P_38_MAPK and diminished intracellular parasite count which again reversed with P_38_MAPK inhibitor application. This acquired evidence confirmed the close association of P_38_MAPK activation and leishmanial control. The same observation pattern was elaborated for NF-κBp65 (Fig. 2**d** and **Supplementary Figure** S3B). Again, the application of inhibitor compromised Sp A mediated intracellular parasite control with associated NF-κBp65 down-expression. Impaired P_38_MAPK expression during host-parasite interaction at the primary hours proven beneficial during subsequent *L. donovani* infection establishment (24) and P_38_MAPK inactivation by *Leishmania* parasites can lead to the iNOS inhibition and inferior NO production (38).

Host invasion by intra-macrophage pathogens is frequently associated with the restraining of NF-κBp65 levels in infected macrophages, and this reduction can modulate the gene expressions necessary during the normal immunological clearance of the intracellular parasites (54). Phosphorylated P_38_MAPK also promotes the association of NADPH oxidase at the cell membrane and resulted in the production and release of superoxide (O_2_^-^) (55). Enhanced NF-κBp65 activation leads to the ensuing transcription of inducible nitric oxide synthase (iNOS) and production of nitric oxide (56) as well as the production of tumour necrosis factor (TNF) and IL-6 (2). Sp A was monitored to activate Gp91-phox, a glycosylated subunit of NADPH oxidase flavocytochrome b558 and also a marker of the NOX activity (57) with subsequent upregulated p-P_38_MAPK and p-NF-κBp65 expression (Fig. 3**b** and **Supplementary Figure S5**). Expression and upregulated nuclear translocation of NF-κBp65 and c-JUN, is required for the enhanced production of pro-inflammatory cytokines, and this event is simultaneously associated with decreased translocation of NF-κBp50 for the ensured functional activity of NF-κB, the key factor regulating the expression of TNF-α (tumor necrosis factor-α), IL-12 and iNOS all are essential for intracellular leishmanial control (58, 5).

Chemotherapeutic agents e.g., miltefosine can upregulate intracellular ROS and RNS and can initiate apoptosis-like cell death in intracellular Leishmania parasites which are widely accepted mechanisms of anti-leishmanial activity of miltefosine (59). It is also reported that, during initial internalization of *Leishmania* inside macrophages, O2̅ is produced as a result of the phagocytosis oxidative burst (59), afterward, the second oxidant NO is generated when the macrophage got activated by Interferon-γ and TNF-α (60, 61) both of them is subverted during leishmaniasis infection or infected cells become less responsive (62, 5). Importantly, for their persistence inside macrophages, *Leishmania* parasites thus implement strategies to overcome the oxidative stress by the cells, and thus the infected cells are having a lower presence of O2̅ or NO as a part of survival strategies (59,61,5). Improved production of ROS after P2X_7_R activation is well recognized in MΦs (63), mediated byNOX2 which additionally can also augment mitochondrial ROS (64). *Leishmania* parasites inside macrophages acclimatize various intimidating conditions during the increment of ROS and RNS by host macrophages (65). At the onset of leishmanial infection, superoxide production resulted from the oxidative burst due to phagocytosis (66) and later NO is produced after activation of macrophages by IFN-γ and TNF(67) resulting in the abolition of intracellular parasites (68). The active subversion of ROS and NO production due to leishmanial infection was noticed to be minimized during Sp A treatment and application of their abrogators minimized the anti-leishmanial effect of the compound (Fig. 3**a** and **Supplementary Figure** S4A, S4B) and thus confirmed the importance of ROS and NO in RAW 264.7 intracellular parasite removal.

P2X_7_R activation mediates large-scale ATP release and forms a membrane pore or prompts downstream purinergic signaling. One major effect is the inflammasome activation through maturation and release of pro-inflammatory cytokines, e.g., IL-1β and IL-18, increment of ROS, and RNS (47). In respect to the intra-macrophage pathogens, the activation of M1 macrophages can subsequently release various cytokines, like interferon-gamma (IFN-γ) and tumor necrosis factor-alpha (TNF-α) can eliminate them by triggering an oxidative burst. Conversely, M2 activation is beneficial for the intra-macrophage pathogens, like *Leishmania* as M2 macrophages do benefit parasite survival in the infected macrophages and disease progression (69). P2X_7_R activation in M1 macrophages was reported to activate the NLRP3 inflammasome, and release of IL-1β and IL-18 along with other inflammatory proteins like lysosomal proteases (cathepsin B and S) or inflammasome components (ASC, NALP3, caspase-1) (70) all of them are useful in intracellular pathogen removal. Reports are stating that P2X_7_ KO mice were more prone to *L. amazonensis* infection with increased lesion and parasitic load (45, 47).

Meanwhile, a molecular docking study was performed to get a primary idea about the binding interaction between Sp A and its putative target P2X_7_. Few hydrogen bonding interactions were observed between Sp A and amino acids at the allosteric site binding of P2X_7_ (PDB: 5U1X). The docked conformation of Sp A with P2X_7_ indicates the non-sugar part of Sp A is exposed to the solvent-accessible surface area.

Finally, *in vitro* results from activity evaluation of Sp A mediated intracellular anti-leishmanial effect were directed to the inference that P2X_7_R-P_38_MAPK activation is the key mediator of the said action with the downstream involvement of Ca^++^ regulated ROS, NO, and phagolysosomal fusion as the effecter cascade. A flow diagram of which was depicted in Fig. 6**d**. Additionally, *in vivo* evaluation in determining the effect of Sp A also established the fact that P2X_7_R and P_38_MAPK expression was upregulated (Fig. 5b, c) and confirmed the *in vitro* findings.

## 4. Methods

### 4.1. Materials

Cell culture medium, Roswell Park Memorial Institute-1640(RPMI-1640) and M-199, foetal bovine serum (FBS), penicillin-streptomycin (PS), and HEPES were acquired from Gibco BRL, Grand Island, NY, USA. Tissue culture plastic wares were acquired from NUNC (Roskilde, Denmark) and DAPI (4*′*,6-diamidino-2-phenylindole dihydrochloride), Lysotracker Red, CFSE (Carboxyfluoresceinsuccinimidyl ester), and FURA 2-AM (Fura-2-acetoxymethyl ester) from Invitrogen, CA. Red Blood Cell (RBC) Lysis Buffer purchased from Abcam, Cambridge, MA, USA. Lipopolysaccharide (LPS, 0111:B4), was purchased from Sigma Chemical Company, St. Louis, Mo, USA. Primary antibodies, secondary antibodies, and inhibitors were obtained from Santa Cruz Biotechnology, Santa Cruz, CA, and Cell signaling technologies, USA.

### 4.2. Ethical approval

All pathogen and animal-related works were performed according to prior approval from Institutional Biosafety Committee, CSIR-Indian Institute of Chemical Biology (CSIR IICB/IBSC/CERT-32/18-19) and CSIR-IICB-Animal Ethics Committee (IICB/AEC/Meeting/Oct/ 2017), respectively. The care and maintenance of animals were done in agreement with the Committee for the Purpose of Control and Supervision of Experiments on Animals (CPCSEA), guidelines.

### 4.3. Macrophages

Murine macrophage (MΦ) cell line RAW 264.7 (American Type Culture Collection, Rockville, MD) was cultured in Roswell Park Memorial Institute medium 1640 (RPMI-1640), supplemented with 10% heat-inactivated FBS and 100 μg/mL of antibiotic (penicillin-streptomycin).

### 4.4. Parasites

Pathogenic *Leishmania donovani* [MHOM/IN/1983/AG83] infection was maintained by the passage in BALB/c mice. *L. donovani* promastigotes, attained from infected BALB/c mice spleen were cultured in complete M199 media (25 mM HEPES, sodium bicarbonate (2.2g/L), 10% FBS and 1% Pen Strep (pH - 7.2-7.4) at 22°C. All pathogen and animal-related works were performed according to prior approval from Institutional Biosafety Committee, CSIR-Indian Institute of Chemical Biology (CSIR IICB/IBSC/CERT-32/18-19) and CSIR-IICB-Animal Ethics Committee (IICB/AEC/Meeting/Oct/2017), respectively. The care and maintenance of animals were done in agreement with the Committee for the Purpose of Control and Supervision of Experiments on Animals (CPCSEA), guidelines.

### 4.5. Preparation and characterization of Spergulin-A

Isolation, characterization of Sp A from the aerial part of a perennial shrub *Glinus oppositifolius* had been described elaborately in our previously reported article (5). Sp A used in this work has been harvested following the same criteria. Briefly, the aerial part of the plant had been dried and subjected to extraction with MeOH at room temperature. The extract was then filtered and the solvent was evaporated using a rotary evaporator and then lyophilized to obtain the crude MeOH extract. The extract was then suspended in Milli-Q H_2_O and partitioned with EtOAc and *n*-BuOH.

The *n*-BuOH fraction was then chromatographed on silica gel using a step gradient, to get a total of five to six fractions. Repeated silica gel column chromatography was performed for the isolation of compounds. Further chromatography was performed over a silica gel column and eluted with increasing polarity of CHCl_3_– MeOH for achieving further diversified fractions (20 to 30). Among these, suggested on having similar pattern the HPLC analyses with four major peaks some fractions were mixed (Discovery® RP amide C16, 5 µm, 25 cm× 4.6 mm, eluted with MeOH–H_2_O 60:40, at 1 mLmin^−1^, ELS-detector). The mixed fractions were purified using Prep-HPLC. All the peaks were then collected and evaporated under reduced pressure using a rotary evaporator at 45°C. Sp A was also characterized as mentioned earlier (5) using ESI Mass spectra, 1H and 13C NMR, Diaion HP 20 was used for column chromatography, silica gel (60 F254) was used for TLC. The final yield was a whitish, amorphous powder-like component with a final yield of 7mg/kg of the dried *G. oppositifolius*. The stock solution of Sp A has been prepared in PBS (Phosphate-buffered saline) at 2 mg/mL concentration.

### 4.6. Parasite infection and treatment

A ratio of 1:10 (MΦs: parasite) *L. donovani* promastigotes was used to infect RAW 264.7MΦs. RAW 264.7MΦs were incubated with parasites for 4 h, washed with PBS, and either fresh media was added or treated with inhibitors for the required time. To study the anti-leishmanial activity, normal and infected RAW 264.7MΦs were treated with Sp A at effective concentrations up to 24h. Miltefosine was used as a positive anti-leishmanial reference. All the measures are taken not to contaminate the experimental setup with endotoxin that can result in the falsified interpretation of experimental data. Additionally, LPS (1 EU/mL) were used for 24 h with control RAW 264.7MΦs as a positive reference to evaluate total P_38_MAPK and p-P_38_MAPK during treatment.

### 4.7. FACS and Confocal microscopy with CFSE tagged L. donovani

RAW 264.7MΦs were parasitized with CFSE-tagged promastigotes (25μM/1×10^6^ promastigotes in 1 mL media for 30 min) and subjected to treatment with Sp A(10, 20, 30μg/mL)for 24 h.

For confocal microscopy, RAW 264.7MΦs (1×10^5^/glass coverslips) were washed twice with PBS after infection and treatment, then fixed in chilled 70% ethanol. The nuclei were stained with DAPI and observed under OlympusFluoview FV10i confocal microscope with a 60X objective lens. For FACS, MФs (1×10^5^/well in a six-well plate) were rinsed twice with PBS after infection and treatment, harvested by scraping, resuspended in 400 μL PBS, and analyzed by BD FACS LSR Fortessa with excitation at 494 nm and emission at 518 nm.

### 4.8. Measurement of Phospho-P_38_MAPK, Gp91-phox, Phospho-NF-κBp65

For phospho-P_38_MAPK and phospho-NF-κBp65 determination control, infected, and treated RAW 264.7MΦs after completion of the treatment schedule of 24 h, were washed with PBS, then fixed with 70% ethanol, after perforation with Triton-x 100 solution (0.5%) labeled with primary and respective fluorochrome conjugated antibodies. 20µM of SB203580 was used to inhibit p-P_38_MAPK and 20µM of BAY-11-7082was used as a p-NF-κBp65 inhibitor. The inhibitors at mentioned concentrations were provided to the specific panels for 3 h and then washed and proceeded for Sp A treatment as scheduled for 24 h. For the measurement of Gp91-phox, the above procedures were followed with specific primary and secondary antibodies. Cells were then analyzed in BD FACS LSR Fortessa.

### 4.9. Measurement of P2X_7_R with BB-G

To determine P2X_7_R level, control, infected, and treated RAW 264.7MΦs were washed with PBS, then fixed with 70% ethanol, after perforation with Triton-x 100 solution (0.5%) labeled with primary and respective fluorochrome conjugated antibodies. Before treatment with Sp A, cells were treated with 10µM of Brilliant Blue G (BB-G) to inhibit P2X_7_R in macrophage cells and present for 24 h of the incubation period.

### 4.10. Confocal Microscopy

After fixation with 70% ethanol and perforation, RAW 264.7MΦs were stained with primary and fluorochrome conjugated secondary antibodies. Then slides were mounted and nuclei were stained with DAPI in case of translocation of p-P_38_MAPK, P2X_7_R, and Gp91-phox.To study the leishmanicidal activity, CFSE was used to stain *L. donovani* and Lysotracker was used to monitor acidic organelles in cells. Rab5 and CathepsinD were stained using primary and respective fluorochrome conjugated secondary antibodies. All specimens were observed under an Olympus Fluoview FV10i confocal microscope with a 60X objective lens.

### 4.11. Measurement of calcium release

Treated and untreated RAW 264.7MΦs were collected, after washing twice with PBS, treated with FURA 2 AM (Molecular Probes) at 37°C. After 45 min of incubation, the cells were washed with PBS, resuspended in fresh PBS, and examined by BD FACS LSR Fortessa with an excitation wavelength of 380 nm and an emission wavelength of 510 nm.

### 4.12. Confocal microscopy analyses of intracellular ROS and NO

For confocal microscopy analyses of intracellular ROS and NO, RAW 264.7MΦs (1×10^5^/glass coverslips) were washed twice with PBS after infection and treatment and exposure to fluorescence probes, briefly, cells were incubated in PBS containing 7.0 μM DAF-2DA (at 37°C for 30 min) or 20 μM DCF-DA (at 37°C for 15 min) then fixed in chilled 70% ethanol. The nuclei were stained with DAPI and observed under OlympusFluoview FV10i confocal microscope with a 60X objective lens. NAC at the concentration of 2 mM and NMMA at the concentration of 100µMwas used as ROS and NO abrogators respectively.

### 4.13. Western blot analysis

40 µg of proteins harvested from control, *L. donovani* infected, Sp A (30µg/mL), LPS (1 EU), P_38_MAPK inhibitor and P2X_7_R inhibitor exposed RAW 264.7MΦs were electrophoretically separated in SDS-PAGE and then transferred to PVDF membrane, blocked with BSA and incubated with respective primary antibodies overnight. The membranes were then incubated with HRP conjugated secondary antibodies, and immunoreactive bands were visualized by adding proper substrates. β-Actin was used as a loading control (5). p-P_38_MAPK and p-NF-κBp65 expression had been normalized with total P_38_MAPK and NF-κBp65 expression respectively.

### 4.14. In vivo leishmanicidal activity

Young healthy male BALB/c mice of 4 weeks of age and 18±2 gm of weight were given *L. donovani* [MHOM/IN/1983/AG83] promastigote infection (1×10^7^ parasite in 100 µL sterile saline inside the heart of the mice) within 5^th^ *in vitro* culture passage. Confirmation of the *L. donovani* infection in the infected BALB/c was done after 2 weeks and thereafter given an adequate time (30 days) for the infection propagation in vital organs like the liver, spleen, and bone marrow. Male BALB/c mice, therefore, attained 8 weeks of age during treatment and weighted 26±2 gm were included in further study. Infected mice were then arbitrarily divided into two groups; one infected control group received only the vehicle and one Sp A treated group that received Sp A at 30 mg/kg b.wt. of the mice, intraperitoneally in an emulsified saline solution of 1 mg/mL concentration for every alternate day for 10 doses. Besides, one un-infected control group with male BALB/c mice (8 weeks of age, weight 26±2 gm) was formed that also received Sp A at 30mg/kg b.wt. for every alternate day for 10 doses. Each group of mice was formed with 7±2 animals. Mice were sacrificed on the 20^th^ day after starting the treatment, i.e., approximately attained 11^th^ week of age. There was no visible change or difference of appearance, fur quality, movement, temper, food and water uptake, excretion, or urination of the experimental mice of all three groups.

Mice after treatment schedule were euthanatized by Ketamine HCl (22-24 mg/kg i/m) under strict guidelines approved by the CSIR-IICB-Animal Ethics Committee (IICB/AEC/Meeting/Oct/ 2017) and in agreement with the CPCSEA guidelines. All the primary cell isolation had been performed under a sterile condition with sterile equipment and media.

For isolating bone-marrow-derived macrophages, femurs, and tibia from the mice had been cleanly isolated, bone marrow was rinsed out in a sterile 15 mL centrifuge tube with DMEM (Dulbecco’s Modified Eagle Medium) through the little hole (opening of the medullary cavity). The initial cell clump was broken down thereafter the cell suspension was passed through 40 µm cell strainer, and harvested by centrifugation spin at 2000 rpm for 5 min. RBC was then lysed by RBC lysis buffer (Abcam) for 5-10 minutes at room temperature and again centrifuged at 400g for 5 minutes and the cell pellet had been re-suspended in complete DMEM. Cells were then placed in a sterile Petri plate for 6 h in a CO2 incubator and only the adherent cells were considered for further analysis.

Spleen macrophages were harvested from the same mice used to collect bone-marrow-derived macrophages. Spleens were excised from the euthanatized mice under sterile conditions and spliced with forceps. The spliced spleens were then agitated for 15 min at 37°C with collagenase A (0.163 U/ml in HBSS) while the tissue splices were just submerged and a single cell suspension was prepared by carefully passing the spliced spleen fragments through a 100 µm Falcon nylon cell strainer (Corning, NY, USA) and cells were re-suspended in complete DMEM media, RBC then lysed by RBC lysis buffer as mentioned earlier and been resuspended in complete DMEM. Cells were then placed in a sterile Petri plate for 6 h in a CO2 incubator and only the adherent cells were considered for further analysis.

For phospho-P38 MAPK, phospho-NF-κBp65, and P2X_7_R protein determination, control and treated primary cells were stained using primary and fluorochrome conjugated secondary antibodies. Then Cells were then analyzed by BD FACS LSR Fortessa with respective excitation and emission wavelength.

### 4.15. In silico analysis

A molecular docking study (71–73) was employed to check the binding mode of the investigated compound toward its putative target P2X_7_R. Chem 3D ultra 8.0.3 (74) software was used to construct the three-dimensional structure of the investigated compound (74). The energy minimization of the compound was done by a molecular orbital package (MOPAC) module with 100 iterations and a minimum RMS gradient of 0.10 (72). The preparation of the ligand structure prior to docking is performed by incorporating polar hydrogen atoms along with Gasteiger partial charges. The torsions are allowed to rotate during docking (75).

The crystallographic structures of P2X7R (PDB: 5U1X) were used for this analysis (76). The macromolecules were prepared by using Discovery Studio 3.0 (77). Polar hydrogen atoms were added to the protein by using the AutoDock Tools (ADT) (78). The grid box was selected 5U1X centered at the corresponding ligand crystal structure by maintaining a spacing of 1 Å (72). Finally, the docking was performed by using the AutoDock Vina (79). The docked poses were visualized in Discovery Studio 3.0 (77).

### 4.16. Statistical analysis

All experimental values were denoted as mean ± SEM acquired from at least three replicate experiments. Statistical significance and variations among groups were evaluated with One-Way analysis of variance (ANOVA) followed by Dunnett’s test. P values ≤ 0.05 (* or **#**) or ≤ 0.01 (** or **##**) were considered indicative of significance.

## Authorship Contribution

**Niladri Mukherjee:** Conceptualization, Methodology, Data Curation, Formal analysis, Funding acquisition, Investigation, Project administration, Validation, Visualization, Data curation, Writing - original draft. **Saswati Banerjee:** Conceptualization, Data Curation, Investigation, Methodology, Validation, Writing - original draft. **Sk. Abdul Amin:** Methodology, Data Curation, Formal analysis, Software, Validation, Writing - original draft. **Tarun Jha:** Conceptualization, Data Curation, Formal analysis, Resources, Software, Supervision. **Sriparna Datta:** Conceptualization, Formal analysis, Investigation, Project administration, Resources, Supervision. **Krishna Das Saha:** Conceptualization, Data Curation, Formal analysis, Funding acquisition, Investigation, Project administration, Resources, Software, Supervision, Validation, Visualization.

## Declaration of Competing Interest

The authors declare that there is no conflict of interest.

## Funding

This research was funded by the Science and Engineering Research Board (SERB), Govt. of India (GrantNo.<u>PDF/2016/001437</u> and <u>EMR/2015/001674</u>).

## Acknowledgments

The manuscript has been critically checked by the full version of the Grammarly software and Urkund for language corrections and plagiarism.

## Availability of data and materials

All data generated or analysed during this study are included in this article (and its Supplementary Information files). Requests for materials should be made to the corresponding author.

## Appendix A. Supplementary material

Supplementary data to this article can be found online at https://doi.org/xxxx/xxxxxxx.xxxx.xxxx.

**Supplementary Methods:** FACS analyses of intracellular ROS and NO

**Supplementary Figure 1 (S1):** (S1A) FACS analyses of CFSE-tagged *L. donovani* and subsequent P2X_7_R expression. (S1B) Sp A mediated anti-leishmanial activity and subsequent aversion by BB-G (number of parasites/20 MФs)

**Supplementary Figure 2 (S2):** (S2A) Mean fluorescence index values of the expressions of both P2X_7_R and Ca^++^. (S2B) P2X_7_R and Ca^++^ expressing RAW 264.7 MΦ cells against the parent population during parasite infection and Sp A treatment.

**Supplementary Figure 3 (S3):** (S3A) Flow cytometric analyses of *p-*P_38_MAPK expression alteration. (S3B) Flow cytometric analyses of NF-κBp65 expression alteration.

**Supplementary Figure 4 (S4):** (S4A) Evaluation of changes in ROS production. (S4B) Intracellular NO production alterations. (S4C) Dose-dependent evaluation of extracellular NO production.

**Supplementary Figure 5 (S5):** Confocal microscopy of co-expression analyses of Gp91-phox and p-P_38_MAPK during infection and Sp A treatment.

**Supplementary Figure 6 (S6):** Representative of gating of primary cells evaluated for expression analyses during FACS.

## Notes

### Competing Interest Statement

The authors have declared no competing interest.

### Summary of Updates

Little has been changed to support the results and discussion more efficiently

## References

1. Tiuman TS, Santos AO, Ueda-Nakamura T, Filho BP, Nakamura CV (2011) Recent advances in leishmaniasis treatment. Int J Infect Dis 15: e525–e532.

2. Banerjee S, Bose D, Chatterjee N, Das S, Chakraborty S, Das T, Saha KD (2016) Attenuated *Leishmania* induce pro-inflammatory mediators and influence leishmanicidal activity by p38 MAPK dependent phagosome maturation in *Leishmania donovani* co-infected macrophages. Sci Rep 6: 22335.

3. Alvar J, Vélez ID, Bern C, Herrero M, Desjeux P, Cano J, Jannin J, den Boer M (2012) Leishmaniasis worldwide and global estimates of its incidence. PLoS One 7: e35671.

4. Freitas EO, Leoratti FMS, Freire-de-Lima CG, Morrot A, Feijó DF (2016) The contribution of immune evasive mechanisms to parasite persistence in visceral leishmaniasis. Front Immunol 7: 153.

5. Banerjee S, Mukherjee N, Gajbhiye RL, Mishra S, Jaisankar P, Datta S, Das Saha K (2019) Intracellular anti-leishmanial effect of Spergulin-A, a triterpenoid saponin of *Glinus oppositifolius*. Infect Drug Resist 12: 2933–2942.

6. Oryan A (2015) Plant-derived compounds in treatment of leishmaniasis. Iran J Vet Res 16:1– 19.

7. Kaur H, Thakur A, Kaur S (2018) Immunoprophylactic potential of a cocktail of three low molecular weight antigens of *Leishmania donovani* along with various adjuvants against experimental visceral leishmaniasis. Iran J Parasitol 13: 11–23

8. Mandal D, Panda N, Kumar S, Banerjee S, Mandal NB, Sahu NP (2006) A triterpenoid saponin possessing antileishmanial activity from the leaves of *Careya arborea*. Phytochemistry 67: 183–190.

9. Delmas F, Di Giorgio C, Elias R, Gasquet M, Azas N, Mshvildadze V, Dekanosidze G, Kemertelidze E, Timon-David P (2000) Antileishmanial activity of three saponins isolated from ivy, a-hederin, b-hederin and hederacolchiside A1, as compared to their action on mammalian cells cultured in vitro. Planta Med 66: 343–347.

10. Maes L, Vanden Berghe D, Germonprez N, Quirijnen L, Cos P, De Kimpe N, Van Puyvelde L (2004) In vitro and in vivo activities of a triterpenoid saponin extract (PX-6518) from the plant *Maesabalansae* against visceral leishmania species. Antimicrob Agents Chemother 48: 130– 136.

11. Miller CM, Boulter NR, Fuller SJ, Zakrzewski AM, Lees MP, Saunders BM, Wiley JS, Smith NC (2011) The Role of the P2X7 Receptor in Infectious Diseases. PloS Pathog 7: e1002212.

12. Aga E, Katschinski DM, van Zandbergen G, Laufs H, Hansen B, Müller K, Solbach W, Laskay T (2002) Inhibition of the spontaneous apoptosis of neutrophil granulocytes by the intracellular parasite *Leishmania major*. J Immunol 169: 898–905.

13. Lees MP, Fuller SJ, McLeod R, Boulter NR, Miller CM, Zakrzewski AM, Mui EJ, Witola WH, Coyne JJ, Hargrave AC (2010) P2X7 receptor-mediated killing of an intracellular parasite, *Toxoplasma gondii* by human and mouse macrophages. J Immunol 184: 7040–7046.

14. Eskra L, Mathison A, Splitter G (2003) Microarray analysis of mRNA levels from RAW 264.7 macrophages infected with *Brucella abortus*. Infect Immun71: 1125–1133.

15. Liu J, Enomoto S, Lancto CA, Abrahamsen MS, Rutherford MS (2008) Inhibition of apoptosis in *Cryptosporidium parvum*-infected intestinal epithelial cells is dependent on surviving. Infect Immun 76: 3784–3792.

16. Del Aguila C, Izquierdo F, Granja AG, Hurtado C, Fenoy S, Fresno M, Revilla Y (2006) *Encephalitozoon microsporidia* modulates p53-mediated apoptosis in infected cells. Int J Parasitol 36: 869–876.

17. Chen S, Ma Q, Krafft PR, Chen Y, Tang J, Zhang J, Zhang JH (2013) P2X7 receptor antagonism inhibits p38 MAPK activation and ameliorates neuronal apoptosis after subarachnoid hemorrhage in the rat. Crit Care Med 41: e466–74.

18. Pfeiffer ZA, Guerra AN, Hill LM, Gavala ML, Prabhu U, Aga M, Hall DJ, Bertics PJ (2007) Nucleotide receptor signalling in murine macrophages is linked to reactive oxygen species generation. Free Radic Biol Med 42: 1506–1516.

19. Gavala ML, Pfeiffer ZA, Bertics PJ (2008) The nucleotide receptor P2RX7 mediates ATP-induced CREB activation in human and murine monocytic cells. J Leukoc Biol 84: 1159–1171.

20. Lee SH, Eom SH, Yoon NY, Kim MM, Li YX, Ha SK, Kim SK (2016) Fucofuroeckol-A from *Eisenia bicyclis* inhibits inflammation in lipopolysaccharide-induced mouse macrophages via downregulation of the MAPK/NF-κB Signaling Pathway. J Chem: 6509212.

21. Cavalli V, Vilbois F, Corti M, Marcote MJ, Tamura K, Karin M, Arkinstall S, Gruenberg J (2001) The stress induced MAP kinase p38 regulates endocytic trafficking via GDI:Rab5 complex. Mol Cell7: 421–432.

22. Prive C, Descoteaux A (2000) *Leishmania donovani* promastigotes evade the activation of mitogen-activated protein kinases p38, c-Jun N-terminal kinase, and extracellular signal-regulated kinase-1/2 during infection of naive macrophages. Eur J Immunol 30: 2235–2244.

23. Jeong HJ, Koo HN, Na HJ, Kim MS, Hong SH, Eom JW, Kim KS, Shin TY, Kim HM (2002) Inhibition of TNF-alpha and IL-6 production by Aucubin through blockade of NF-kappaB activation RBL-2H3 mast cells. Cytokine 18: 252–259.

24. Junghae M, Raynes JG (2002) Activation of p38 mitogen-activated protein kinase attenuates *Leishmania donovani* infection in macrophages. Infect Immun 70: 5026–5035.

25. Gutierrez MG (2013) Functional role(s) of phagosomal Rab GTPases. Small GTPases 4: 148– 158.

26. Fischer W, Urban N, Immig K, Franke H, Schaefer M (2014) Natural compounds with P2X7 receptor-modulating properties. Purinergic Signal 10: 313–326.

27. Bartlett R, Stokes L, Sluyter R (2014) The P2X7 receptor channel: recent developments and the use of P2X7 antagonists in models of disease. Pharmacol Rev 66: 638–675.

28. Stokes L, Layhadi JA, Bibic L, Dhuna K, Fountain SJ (2017) P2X4 Receptor Function in the Nervous System and Current Breakthroughs in Pharmacology. Front Pharmacol 8: 291.

29. Helliwell RM, ShioukHuey CO, Dhuna K, Molero JC, Ye JM, Xue CC, Stokes L (2015) Selected ginsenosides of the protopanaxdiol series are novel positive allosteric modulators of P2X7 receptors. Br J Pharmacol 172: 3326–3340.

30. Orioli E, De Marchi E, Giuliani AL, Adinolfi E (2017) P2X7 Receptor Orchestrates Multiple Signalling Pathways Triggering Inflammation, Autophagy and Metabolic/Trophic Responses. Curr Med Chem 24: 2261–2275.

31. Santos JA, Fidalgo-Neto AA, Faria RX, Simões A, Calheiros AS, Bérenger AL, Faria-Neto HC, Figueiredo MR, Frutuoso VS, Alves LA (2011) Effect of *Rheedia longifolia* Leaf Extract and Fractions on the P2X_7_ Receptor *In Vitro*: Novel Antagonists? J Med Food 14: 920–929.

32. Podolak I, Galanty A, Sobolewska D (2010) Saponins as cytotoxic agents: a review. Phytochem Rev 9: 425–474.

33. Peng W, Cotrina ML, Han X, Yu H, Bekar L, Blum L, Takano T, Tian GF, Goldman SA, Nedergaard M (2009) Systemic administration of an antagonist of the ATP-sensitive receptor P2X7 improves recovery after spinal cord injury. ProcNatl Acad Sci USA 106: 12489–12493.

34. Wang XH, Xie X, Luo XG, Shang H, He ZY (2017) Inhibiting purinergic P2X7 receptors with the antagonist brilliant blue G is neuroprotective in an intranigral lipopolysaccharide animal model of Parkinson’s disease. Mol Med Rep 15: 768–776.

35. Soares-Bezerra R J, Ferreira NC, Alberto AV, Bonavita AG, Fidalgo-Neto AA., Calheiros AS, FrutuosoVda S, Alves LA (2015) An Improved Method for P2X7R Antagonist Screening. PLoS ONE 10: e0123089.

36. Namovic MT, Jarvis MF, Donnelly-Roberts D (2012) High throughput functional assays for P2X receptors. Curr Protoc Pharmacol Chapter 9: Unit 9 15.

37. Biswas D, Qureshi OS, Lee WY, Croudace JE, Mura M, Lammas DA (2008) ATP induced autophagy is associated with rapid killing of intracellular mycobacteria within human monocytes/macrophages. BMC Immunol 9:35.

38. Olivier M, Gregory DJ, Forget G (2005)Subversion mechanisms by which *Leishmania* parasites can escape the host immune response: a signaling point of view. Clin Microbiol Rev 18: 293–305

39. Van Assche T, Deschacht M, da Luz RA, Maes L, Cos P (2011) Leishmania-macrophage interactions: insights into the redox biology. Free Radic Biol Med 51: 337–351.

40. Soulat D, Bogdan C (2017) Function of macrophage and parasite phosphatases in Leishmaniasis. Front Immunol 8: 1838.

41. Coutinho-Silva R, Stahl L, Raymond MN, Jungas T, Verbeke P, Burnstock G, Darville T, Ojcius DM (2003) Inhibition of chlamydial infectious activity due to P2X7R-dependent phospholipase D activation. Immunity 19: 403–412.

42. Coutinho-Silva R, Ojcius DM (2012) Role of extracellular nucleotides in the immune response against intracellular bacteria and protozoan parasites. Microbes Infect 14: 1271–1277.

43. Amaral EP, Ribeiro SC, Lanes VR, Almeida FM, de Andrade MR, Bomfim CC, Salles EM, Bortoluci KR, Coutinho-Silva R, Hirata MH. et al. (2014) Pulmonary infection with hypervirulent Mycobacteria reveals a crucial role for the P2X7 receptor in aggressive forms of tuberculosis. PloS Pathog 10: e1004188.

44. Chaves SP, Torres-Santos EC, Marques C, Figliuolo VR, Persechini PM, Coutinho-Silva R, Rossi-Bergmann B (2009) Modulation of P2X7 purinergic receptor in macrophages by *Leishmania amazonensis* and its role in parasite elimination. Microbes Infect 11: 842–849.

45. Figliuolo VR, Chaves SP, Savio LEB, Thorstenberg MLP, Machado Salles É, Takiya CM, D’Império-Lima MR, de Matos Guedes HL, Rossi-Bergmann B, Coutinho-Silva R (2017) The role of the P2X7 receptor in murine cutaneous leishmaniasis: aspects of inflammation and parasite control. Purinergic Signal 13: 143–152.

46. De Marchi E, Orioli, E, Dal BD, Adinolfi E (2016) P2X7 receptor as a therapeutic target. Adv Protein Chem Struct Biol 104: 39–79.

47. Savio LEB, de Andrade Mello P, da Silva CG, Coutinho-Silva R (2018) The P2X7 Receptor in Inflammatory Diseases: Angel or Demon? Front Pharmacol 9:52.

48. Illes P (2020) P2X7 Receptors Amplify CNS Damage in Neurodegenerative Diseases. Int J Mol Sci 21: 5996.

49. Nunes P, Demaurex N (2010) The role of calcium signaling in phagocytosis. J Leukoc Biol 88: 57–68.

50. Pinton P, Giorgi C, Siviero R, Zecchini E, Rizzuto R (2008) Calcium and apoptosis: ER-mitochondria Ca^2+^ transfer in the control of apoptosis. Oncogene 27: 6407–6418.

51. Moradin N, Descoteaux A (2012) *Leishmania* promastigotes: building a safe niche within macrophages. Fron tCell Infect Microbiol 2: 121

52. Swulius MT, Waxham MN (2008) Ca(2+)/calmodulin-dependent protein kinases. Cell Mol Life Sci 65: 2637–2657.

53. Corre I, Paris F, Huot J (2017) The p38 pathway, a major pleiotropic cascade that transduces stress and metastatic signals in endothelial cells. Oncotarget 8: 55684–55714.

54. Calegari-Silva TC, Pereira RM, De-Melo LD, Saraiva EM, Soares DC, Bellio M, Lopes UG (2009) NF-κB-mediated repression of iNOS expression in *Leishmania amazonensis* macrophage infection. Immunol Lett 1 27:19–26.

55. Tarafdar A, Pula G (2018) The role of NADPH oxidases and oxidative stress in neurodegenerative disorders. Int J Mol Sci 19: 3824.

56. McAdam E, Haboubi HN, Forrester G, Eltahir Z, Spencer-Harty S, Davies C, Griffiths AP, Baxter JN, Jenkins GJ (2012) Inducible nitric oxide synthase (iNOS) and nitric oxide (NO) are important mediators of reflux-induced cell signalling in esophageal cells. Carcinogenesis 33: 2035–43.

57. Yu L, Quinn MT, Cross AR, Dinauer MC (1998) Gp91(phox) is the heme binding subunit of the superoxide-generating NADPH oxidase. Proc Natl Acad Sci USA 95: 7993–7998.

58. Sun Z, Andersson R (2002) NF-kappaB activation and inhibition: a review. Shock 18: 99–106.

59. Deep DK, Singh R, Bhandari V, Verma A, Sharma V, Wajid S, Sundar S, Ramesh V, Dujardin, JC, Salotra P (2017) Increased miltefosine tolerance in clinical isolates of *Leishmania donovani* is associated with reduced drug accumulation, increased infectivity and resistance to oxidative stress. PLoS Neg Trop Dis 11: e0005641

60. Carneiro PP, Conceica̦ OJ, Macedo M, Magalhaes V, Carvalho, EM, Bacellar O (2016) The Role of Nitric Oxide and Reactive Oxygen Species in the Killing of *Leishmania braziliensis* by Monocytes from Patients with Cutaneous Leishmaniasis. PLoS One 11: e0148084.

61. Tiwari R, Banerjee S, Tyde D, et al. (2021) Redox-Responsive Nanocapsules for the Spatiotemporal Release of Miltefosine in Lysosome: Protection against *Leishmania*.Bioconjug Chem 32: 245–253.

62. Kima PE, Soong L (2013) Interferon gamma in leishmaniasis. Front Immunol 4: 156.

63. Lenertz LY, Gavala ML, Hill LM, Bertics PJ (2009) Cell signaling via the P2X(7) nucleotide receptor: linkage to ROS production, gene transcription, and receptor trafficking. Purinergic Signal 5: 175–187.

64. Nakahira K, Haspel JA, Rathinam VA, Lee SJ, Dolinay T, Lam HC, Englert JA, Rabinovitch M, Cernadas M, Kim HP, Fitzgerald KA, Ryter SW, Choi AM (2011). Autophagy proteins regulate innate immune responses by inhibiting the release of mitochondrial DNA mediated by the NALP3 inflammasome. Nat Immunol 12: 222–230.

65. Carneiro PP, Conceição J, Macedo M, Magalhães V, Carvalho EM, Bacellar O (2016) The role of nitric oxide and reactive oxygen species in the killing of *Leishmania braziliensis* by monocytes from patients with cutaneous leishmaniasis. PLoS ONE 11: e0148084.

66. Almeida TF, Palma LC, Mendez LC, Noronha-Dutra AA, Veras PS (2012) *Leishmania amazonensis* fails to induce the release of reactive oxygen intermediates by CBA macrophages. Parasite Immunol 34: 492–498.

67. Gantt KR, Goldman TL, McCormick ML, Miller MA, Jeronimo SM, Nascimento ET, Britigan BE, Wilson ME (2001) Oxidative responses of human and murine macrophages during phagocytosis o*f Leishmania chagasi*. J Immunol 167: 893–901.

68. Bogdan C, Rollinghoff M, Diefenbach A (2000) The role of nitric oxide in innate immunity. Immunol Rev 173: 17–26.

69. Tomiotto-Pellissier F, Bortoleti BTDS, Assolini JP, et al. (2018) Macrophage Polarization in Leishmaniasis: Broadening Horizons. Front Immunol 9: 2529

70. de Torre-Minguela C, Barberà-Cremades M, Gómez AI, Martín-Sánchez F, Pelegrín P (2016) Macrophage activation and polarization modify P2X7 receptor secretome influencing the inflammatory process. Sci Rep 6: 22586.

71. Amin SA, Bhattacharya P, Basak S, Gayen S, Nandy A, Saha A (2017a) Pharmacoinformatics study of piperolactam a from Piper beetle root as new lead for nonsteroidal anti fertility drug development. Comput Biol Chem 67: 213–224.

72. Amin SA, Adhikari N, Agrawal RK, Jha T, Gayen S (2017b) Possible Binding Mode Analysis of Pyrazolo-triazole Hybrids as Potential Anticancer Agents through Validated Molecular Docking and 3D-QSAR Modeling Approaches. Lett Drug Des Discov 14: 515–527.

73. Bhardwaj B, Baidya ATK, Amin SA, Adhikari N, Jha T, Gayen S (2019) Insight into structural features of phenyltetrazole derivatives as ABCG2 inhibitors for the treatment of multidrug resistance in cancer. SAR QSAR Environ Res 30: 457–475.

74. Chem 3D Pro Version 8.0 And Chem Draw Ultra Version 8.0 Are Software Programs Developed by Cambridge Soft Corporation. USA.

75. Lv PC, Li HQ, Sun J, Zhou Y, Zhu HL (2010) Synthesis and biological evaluation of pyrazole derivatives containing thiourea skeleton as anticancer agents. Bioorg Med Chem 18: 4606–4614.

76. https://www.rcsb.org/structure/5U1X: https://www.rcsb.org/structure/5U1X (as accessed on 14th August 2019)

77. Discovery Studio 3.0 (DS 3.0), Accelrys Inc., CA, USA, 2015.

78. Autodock Tools 1.5.6 (ADT)/MGL Tools 1.5.6, The Scripps Research Institute, CA, USA, 2012. Available at http://mgltools.scripps.edu.

79. Trott O, Olson AJ (2010) AutoDock Vina: Improving the speed and accuracy of docking with a new scoring function, efficient optimization, and multithreading. J Comput Chem 31: 455–461.

